# Despite of DNA repair ability the Fanconi anemia mutant protein FANCGR22P destabilizes mitochondria and leads to genomic instability via FANCJ helicase

**DOI:** 10.1101/2020.01.15.907303

**Authors:** Jagadeesh Chandra Bose K, Bishwajit Singh Kapoor, Kamal Mondal, Subhrima Ghosh, Raveendra B. Mokhamatam, Sunil K. Manna, Sudit S. Mukhopadhyay

## Abstract

Fanconi anemia (FA) is a unique DNA damage repair pathway. Almost twenty-two genes have been identified which are associated with the FA pathway. Defect in any of those genes causes genomic instability, and the patients bear the mutation become susceptible to cancer. In our earlier work, we have identified that Fanconi anemia protein G (FANCG) protects the mitochondria from oxidative stress. In this report, we have identified eight patients having mutation (C.65G>C; p.Arg22Pro) in the N-terminal of FANCG. The mutant protein hFANCGR22P is able to repair the DNA and able to retain the monoubiquitination of FANCD2 in FANCGR22P/FGR22P cell. However, it lost mitochondrial localization and failed to protect mitochondria from oxidative stress. Mitochondrial instability in the FANCGR22P cell causes the transcriptional down-regulation of mitochondrial iron-sulphur cluster biogenesis protein Frataxin (FXN) and resulting iron deficiency of FA protein FANCJ, an iron-sulphur containing helicase involved in DNA repair.

## Introduction

Nuclear genomic instability is a common phenomenon and prerequisite for cancer. Genomic stability is maintained by the balance between the rate of DNA damage and rate of repair. Non repairable damage of the genome or genomic mutation may cause loss of heterozygosity (LOH), may activate proto-oncogenes, may inactivate tumor suppressor genes and/or can alter the regulation of genes associated with cell cycle and cellular signals(PC, 1976; Philpott et al., 1998; Veatch et al., 2009). These DNA damaging agents are either exogenous or endogenous. The most abundant endogenous DNA damaging agents are oxidative radicals which are produced primarily by mitochondria. Several studies suggest that one of the causes of genomic instability is the overproduction of reactive oxygen species (ROS) resulting from mitochondrial dysfunction (Vives-Bauza et al., 2006). When the extent of irreparable damage is extensive, then the cell undergoes apoptotic death, a normal phenomenon which is controlled by mitochondria (Kujoth et al., 2005). Therefore, the mitochondrion’s role in malignancy is considerable because in addition to critical changes in metabolism mitochondria determine the balance between survival and death. However, the precise mechanism of how mitochondria maintain nuclear genomic stability is not clearly known. Alterations of both the mitochondrial and nuclear genomes have been observed in various types of cancers (Larman et al., 2012). In yeast, an association of mitochondrial function with genomic DNA integrity has been reported(Flury et al., 1976). Recently, it has been shown that under certain conditions, mitochondrial caspase may lead to nuclear DNA damage and genomic instability (Ichim et al., 2015). Daniel E. Gottschling’s group created a specific mutant strain of *S cerevisiae* and showed that genomic stability is maintained by iron-sulphur cluster synthesis, an essential mitochondrial function (Veatch et al., 2009). Loss of mitochondrial membrane potential causes downregulation of iron-sulfur cluster (ISC) biogenesis, an essential mechanism for the Fe-S domain containing proteins involved in nuclear genomic stability(Kispal et al., 1999). However, this observation requires further studies to identify human pathogenic mutants that might be involved in the process.

In this report, we have explored this hypothesis by describing the mutation of a FA patient subtype G (FANCG). FA is a rare, hereditary, genomic instability and cancer susceptibility syndrome. Congenital disabilities and bone marrow failure are the most prominent features of FA patients. After consecutive bone marrow transplantation (BMT), patients suffer from BMT-associated problems and undergo increased cancer risk, including hematological malignancies and head and neck cancer (Rosenberg et al., 2005). To date, twenty two genes have been identified that associate with FA that are primarily involved in a specific type of DNA damage repair; inter-strand crosslink (ICL) repair. ICL is caused by the exogenous alkylating agents or endogenous metabolites such as formaldehydes and acetaldehydes(Bluteau et al., 2016). Upon damage, out of twenty two, eight proteins (A, B, C, E, F, G, L & M) form a complex which is called the FA core complex(Walden and Deans, 2014). The FA core complex formation initiates the monoubiquitination of both FANCD2 and FANCI, which is called the ID2 complex. The ID2 complex binds the damaged part of the chromatin and in association with other FA proteins and non-FA proteins repair the ICL damage. The repairing complex mostly consists of several exonucleases, endonucleases, helicases, and proteins involved in the DNA damage repair by homologous recombination pathway(Walden and Deans, 2014).

FANCJ is a DEAH superfamily 2 helicase and part of the subfamily of Fe-S cluster-containing helicase-like proteins including XPD, RTEL1, and CHL1(Guo et al., 2016). FANCJ cells are highly sensitive to ICL agents, and mutation studies suggest its association with cancer (Brosh and Cantor, 2014). Many genetic and biochemical studies suggest FANCJ is an ATP dependent helicase which unwinds the duplex DNA or resolves G-quadruplex DNA structures (Guo et al., 2014). Thus, FANCJ has an essential role in ICL damage repair and in maintaining genome stability. Recently, it has been shown that a pathogenic mutation in that iron-sulphur (Fe–S) cluster is essential for helicase activity and iron deficiency results in the loss of helicase activity of the FANCJ but not the ATPase activity (Wu et al., 2010).

The sensitivity of the FA cell to the oxidative stress and several protein-protein interaction studies suggest that FA proteins are also involved in oxidative stress metabolism(Mukhopadhyay et al., 2006). In our earlier studies, we have shown that FA subtype G (FANCG) protein interacts with the mitochondrial protein peroxiredoxin 3 (PRDX3), a member of the peroxidase family and neutralizes the mitochondrial oxidative stress. In FANCG cells PRDX3 is cleaved by calpain protease and loses its peroxidase activity. Elevated oxidative stress alters the mitochondrial structure and loss of mitochondrial membrane potential was observed in the FANCG cells (Mukhopadhyay et al., 2006). These results suggest that FANCG protects the mitochondria from oxidative stress by preventing the PRDX3 from calpain-mediated degradation. Many groups including our own have debated the role of FA proteins in mitochondria (Pagano et al., 2014). In this report, we have identified the N-terminal thirty amino acids, which is unique to humans as the mitochondrial localization signal (MLS) of FANCG. Human mutation studies confirmed both the nuclear and mitochondrial roles of FANCG. The objective of the current study was to identify the defect of FANCJ in mutant cells due to oxidative stress-mediated mitochondrial dysfunction. In conclusion, we showed that specific mutations in the mitochondrial localization signal in FANCG result in mitochondrial dysfunction result in genomic instability.

## Results

### Identification of Mitochondrial Localization Signal (MLS) of human FANCG

In our earlier studies, we have shown that human FANCG protein protects the mitochondrial peroxidase PRDX3 from calpain cleavage and subsequently mitochondria from oxidative stress(Mukhopadhyay et al., 2006). Since FA proteins are known to regulate nuclear DNA damage repair (DDR), this brings up the question of how the FANCG protein migrates to mitochondria. Of the thousands of nuclear proteins that migrate to mitochondria (Backes et al., 2018) some have been shown to have mitochondrial localization signals. However, many of them do not have an identifiable signal peptide. Generally proteins migrate to mitochondria through the interaction of TOM (mitochondrial outer membrane protein) and TIM (mitochondrial inner membrane protein) and some proteins enter with the help of carrier proteins(Nickel et al., 2018). Human FANCG contains a TPR motif (tetratricopeptide repeat) which is known to facilitate protein-protein interaction(Wilson et al., 2010). Initially, we thought that FANCG might interact with some TPR-containing TOM proteins. However, immuno-precipitation (IP) studies did not support this idea (data not shown).

There are several online tools available, which can be used for identification of the signal peptide sequence for protein cellular localization. Similarly, some specific tools are also available for identification of mitochondrial localization signals (MLS) (Bannai et al., 2002). We have utilized all the available tools for identification of the MLS of the FA proteins (Table S1A & B; Supple Fig.S1.A, B and C). The iPSORT analysis predicted thirty amino acids at the N-terminal of human FANCG protein as a mitochondrial localization signal (MLS) or Mitochondrial Targeting Peptide (mTP) (Fig.1A). However, when we analyzed the N-terminal of FANCG of other species, mTP was identified only in human FANCG (Fig.1B & C). For confirmation of this result, the expression of the protein in a mammalian cell line is required. As the N-terminal thirty amino acids were predicted as an MLS, we made several N-terminal deletion FANCG constructs (05DEL, 09DEL, 18DEL, 24DEL, and MLSDEL {30 amino acids}) containing ATG sequences as a start codon (Fig. 1D). All the deletion constructs were sequenced and confirmed to retain an open reading frame. To visualize the FANCG expression in the cell line, the wild-type and the deleted constructs were tagged with GFP at the C-terminal end. The mitochondrial marker Mito-tracker (pDsmito-Red) was used in these co-localization studies. Each deletion construct including a wild type control was transiently expressed along with mito-tracker in HeLa cells. The expression of both the constructs was analyzed by deconvolution microscope (Axio Observer.Z1, Carl Zeiss; Axiovision software). The wild-type FANCG fused with GFP showed perfect co-localization with Mito-tracker in mitochondria of HeLa cells (Fig.1D). However, the deleted constructs showed loss of co-localization with Mito-tracker (Fig.1D). The complete loss of co-localization was observed following deletion of 18, 24 and 30 amino acids (MLSDEL) (Fig.1D). These co-localization studies suggest that the in silico predicted N-terminal thirty amino acids are the mitochondrial targeting Peptide (mTP) which actually determines the mitochondrial localization of human FANCG.

**Fig. 1.**
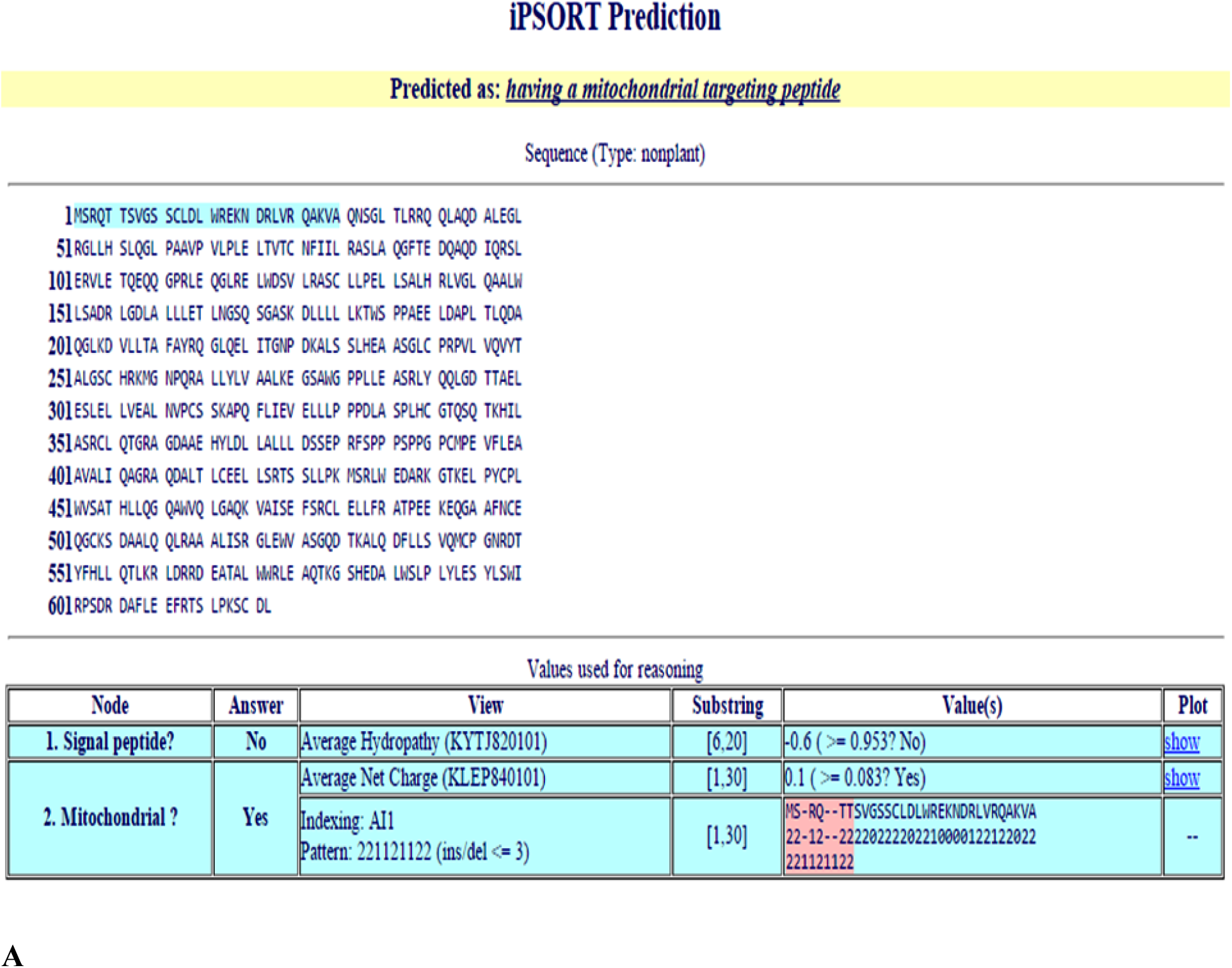
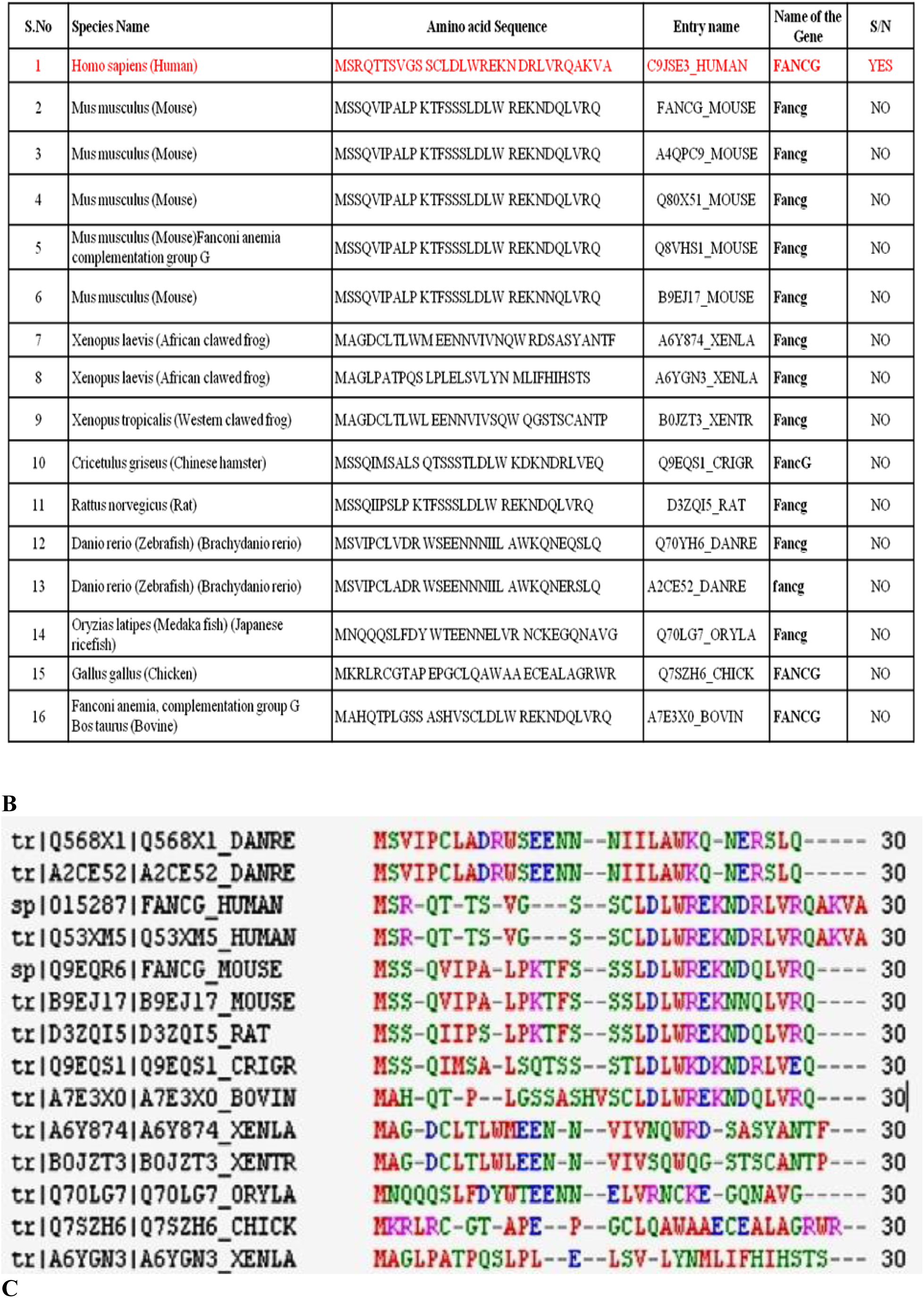
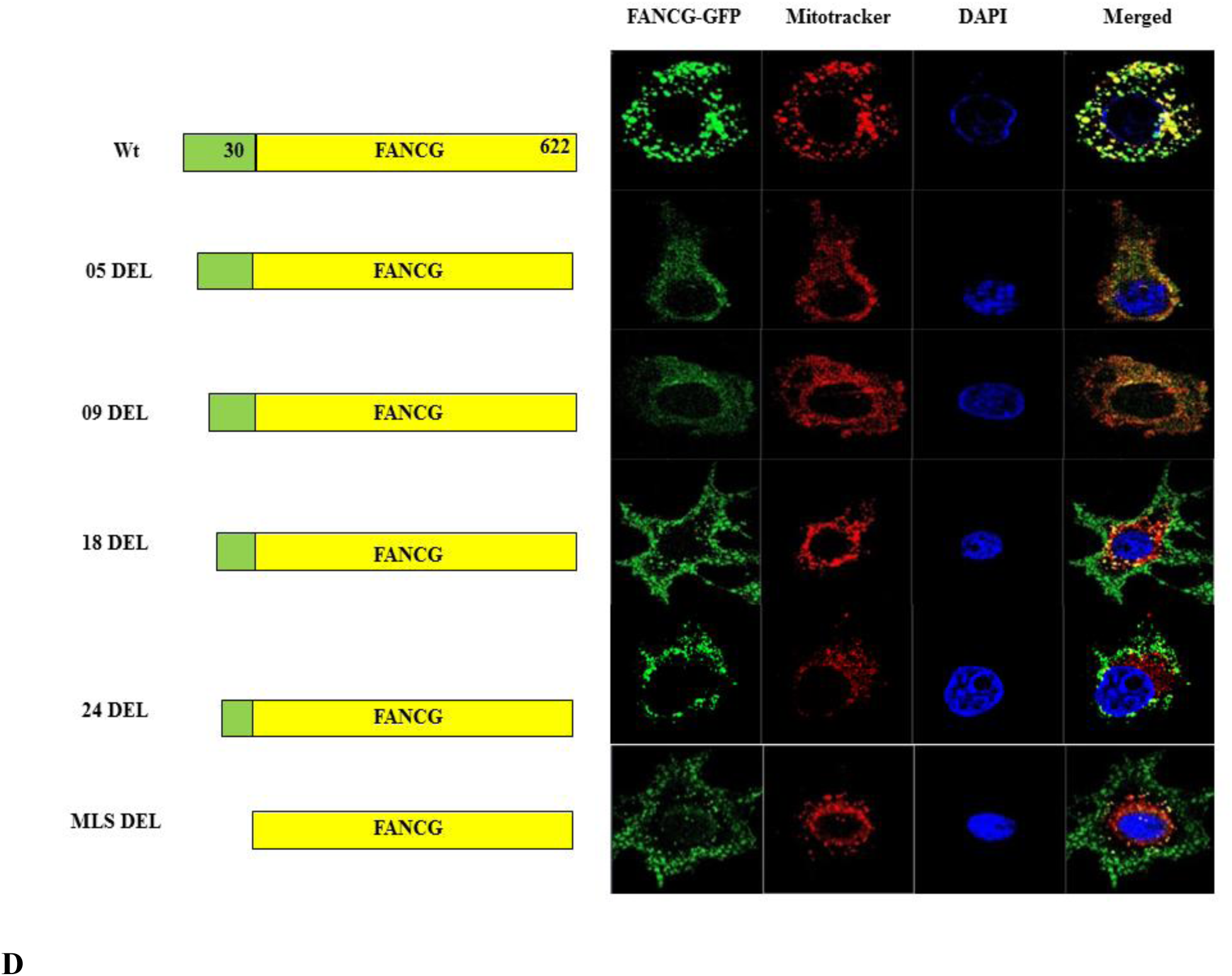
Identification of Mitochondrial Localization Signal of human FANCG. iPsort analysis of the **(A)** human FANCG (highlighted sequences are the predicted as MLS) and **(B)** only N-terminal sequences of FANCG from various species including human.**(C)** Analysis the conserved amino acids among the N-terminal region of the FANCG. **(D)** Co-localization studies of hFANCG-GFP and mitotracker in HeLa cells. Wt= wild type, 05DEL= five amino acids deleted, 09DEL= nine amino acids deleted, 18DEL= eighteen amino acids deleted, 24DEL= twenty four amino acids deleted and MLSDEL= entire MLS deleted. DAPI represents the nucleus.

### Human mutant FANCGR22P lost mitochondrial localization, but not nuclear localization

We further looked for pathogenic mutations in the MLS of FANCG. We identified eight FA patients from the Fanconi Anemia database of Rockefeller University{http://www.rockefeller.edu/fanconi/; **L**eiden **O**pen Source **V**ariation **D**atabase (**LOVD** v.3.0)}. In these eight FA patients due to a single nucleotide change (C.65G>C), the amino acid arginine at the twenty-two position of the MLS was converted into proline (p. (Arg22Pro). The iPSORT analysis predicted the loss of mitochondrial migration of this mutant protein (R22P) (Supple Fig.S2 A & B). We wanted to understand how the single nucleotide change affects the structure of the protein. The crystal structure of human FANCG is not known. In the secondary structure, the MLS of FANCG is made with an alternate stretch of coil and helix. The helix is interrupted by a coil in R22P due to replacement of arginine by proline (Fig.2A). In our modeled structure, similarly, the MLS region of R22P is disrupted due to the replacement of arginine by proline (Fig.2B). How the altered structure affects the mitochondrial migration is not clear. Thus, R22P mutant construct was made and transiently transfected into the HeLa cells along with Mito-tracker. The co-localization study suggested the complete loss of migration of R22P into mitochondria (Fig.2C). Co-localization studies of R22P in different passages of HeLa cells as well as in FANCG parental cells further confirmed its inability of mitochondrial localization (Supple Fig.S2 C & D). We have identified another FANCG patient (S07F) with a mutation in MLS sequence (serine at the seventh position is converted into phenylalanine). However, S07F protein can localize to mitochondria (Fig.2C). FANCG as a member of the FA core complex remains associated with chromatin, and we searched whether the mutant protein R22P can migrate to the nucleus. The cell biology studies in HeLa cells suggest that the mutant protein R22P can migrate to the nucleus upon MMC treatment like the wild-type FANCG (Fig.2D). All these results suggest that FANCG human mutant R22P cannot migrate to mitochondria but can migrate to the nucleus upon MMC treatment.

**Fig. 2.**
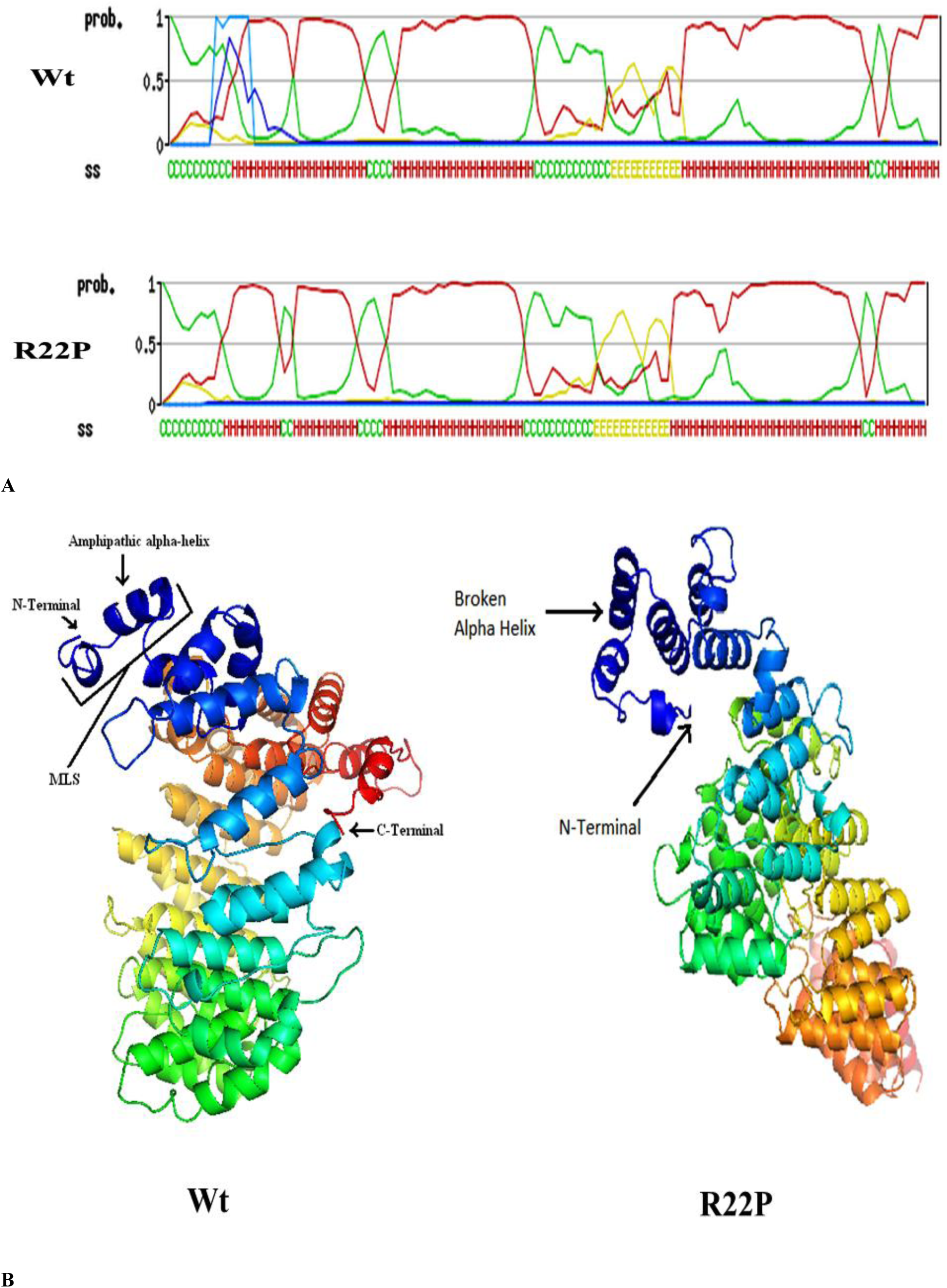
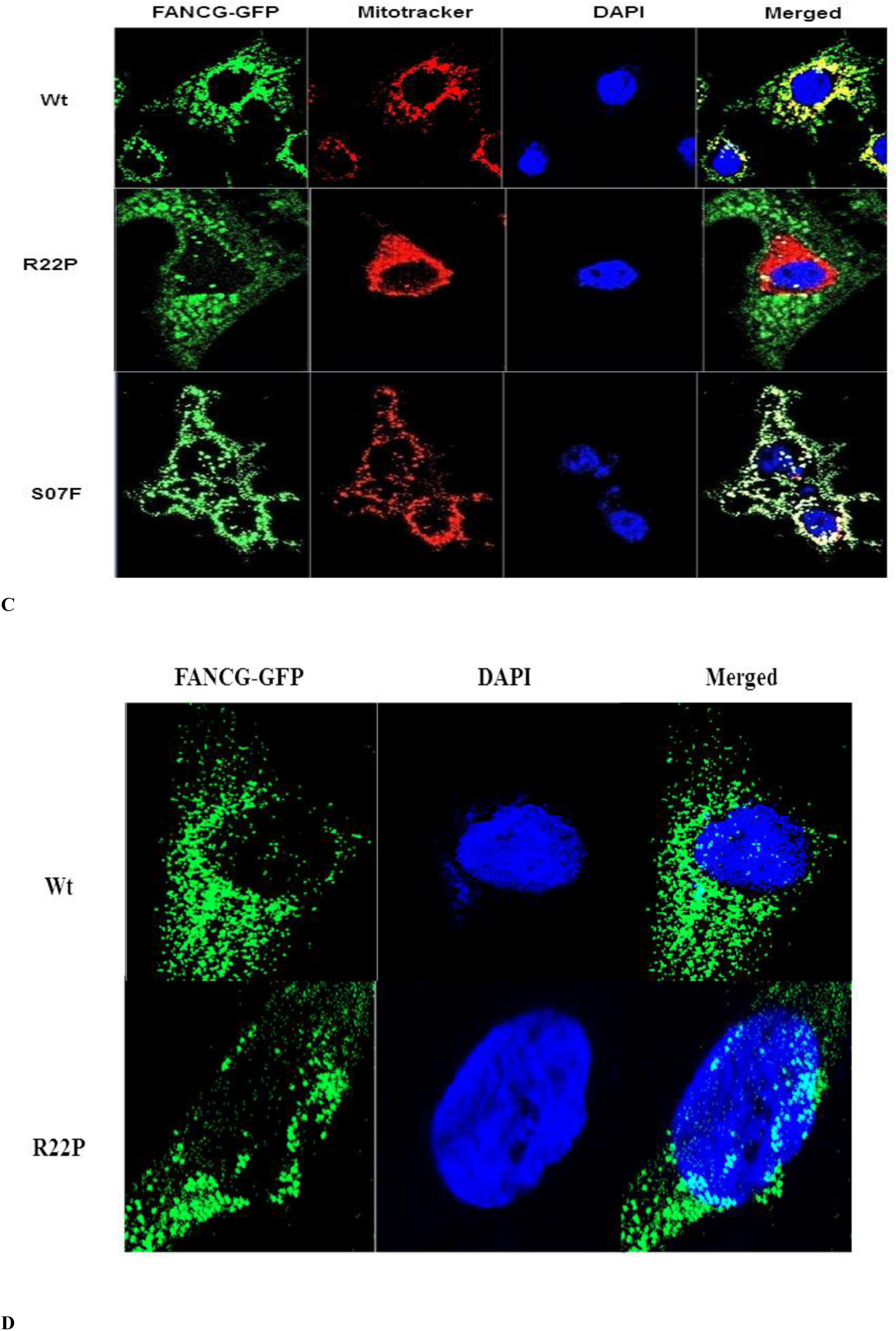
Localization of FANCG R22P in HeLa cells. **(A)** Secondary structure and **(B)** modelled 3D structure of wild type and R22P FANCG. **(C)** Co-localization of wild type (Wt), R22P and S07F of FANCG and mitotracker in HeLa cells. **(D)** Nuclear localization of wt and R22P in HeLa cells treated with MMC.

### FANCGR22P cells are sensitive to oxidative stress but resistant to ICL agents

In order to elucidate whether the FANCG human mutant R22P is functional in the nucleus or not, the R22P stable cell line (Fig.3A & 3B) was developed in the background of FANCG parental cells (lacking FANCG; obtained from Dr. Agata Smogorzewska’s lab, The Rockefeller University, NY) (Fig.3A & 3B). The MMC-mediated FANCD2 monoubiquitination was analyzed in the R22P stable cells, FANCG corrected cells and in FANCG parental cells (Fig.3C). Surprisingly, like FANCG-corrected cells FANCD2 monoubiquitination was observed in the R22P stable cell upon MMC treatment (Fig.3C, lane 1 and 2). Monoubiquitination of FANCD2 is absent in FANCG parental cells treated with MMC (Fig.3C lane 3) and also in the cells not treated with MMC (Fig.3C, lane 4, 5 and 6). This experiment confirmed that FANCG human mutant protein R22P is able to participate in the formation of FA core complex in the nucleus. In order to understand the DNA repair ability of the R22P stable cells, drug sensitivity tests were performed. FANCG corrected, parental and R22P cells were treated with increasing concentration of MMC and cisplatin separately for two and five days. Cell survival was determined both by MTT and Trypan blue assay (Fig 4 & Supple Fig S3). To our surprise, even with five days of treatment with drugs in increasing concentration, the R22P stable cells showed resistance to both MMC and cisplatin like the FANCG corrected cells (Fig.4A, B, C & D). Even with formaldehyde treatment for two hrs, the R22P stable cells showed resistance compared to FANCG parental cells (Fig.4G & Supple Fig.S3G). These drug sensitivity results all suggested that FANCG human mutant protein R22P can form FA core complex and can repair the ICL damage.

**Fig. 3.**
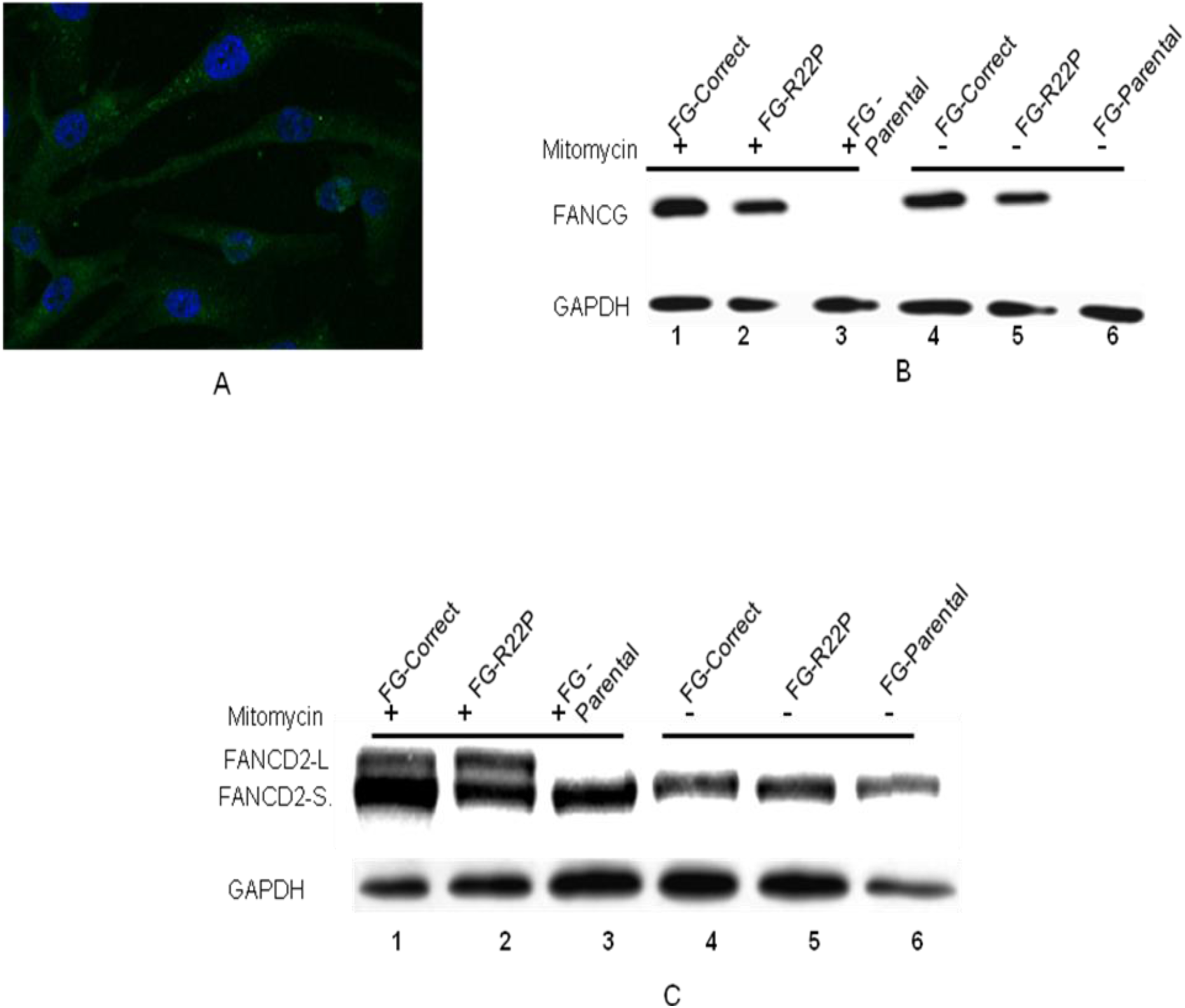
Development of FANCGR22P stable cell line. R22P construct was stably integrated into the genome of FANG parental cell by Lenti vector pLJM1-EGFP (Addgene). (A) GFP expression confirms the stable expression of R22P in FANCG parental cell. (B) Cells were treated with (lane 1, 2 and 3) and without (4,5 and 6) MMC and cell lysates were used for Western blot with FANCG antibody. Expression of FANCGR22P was confirmed in the stable cell (lane 2&5). (C) FANCD2 monoubiquitination studies of the FANCG corrected, FG-R22P and FG-parental cells. Cells were treated with (lane 1.2 and 3) and without (4, 5 and 6) MMC and blotted with FAND2 antibody. FANCD2-L represents the monoubiquitinated and FANCD2-S represents the normal FANCD2 proteins.

**Fig. 4.**
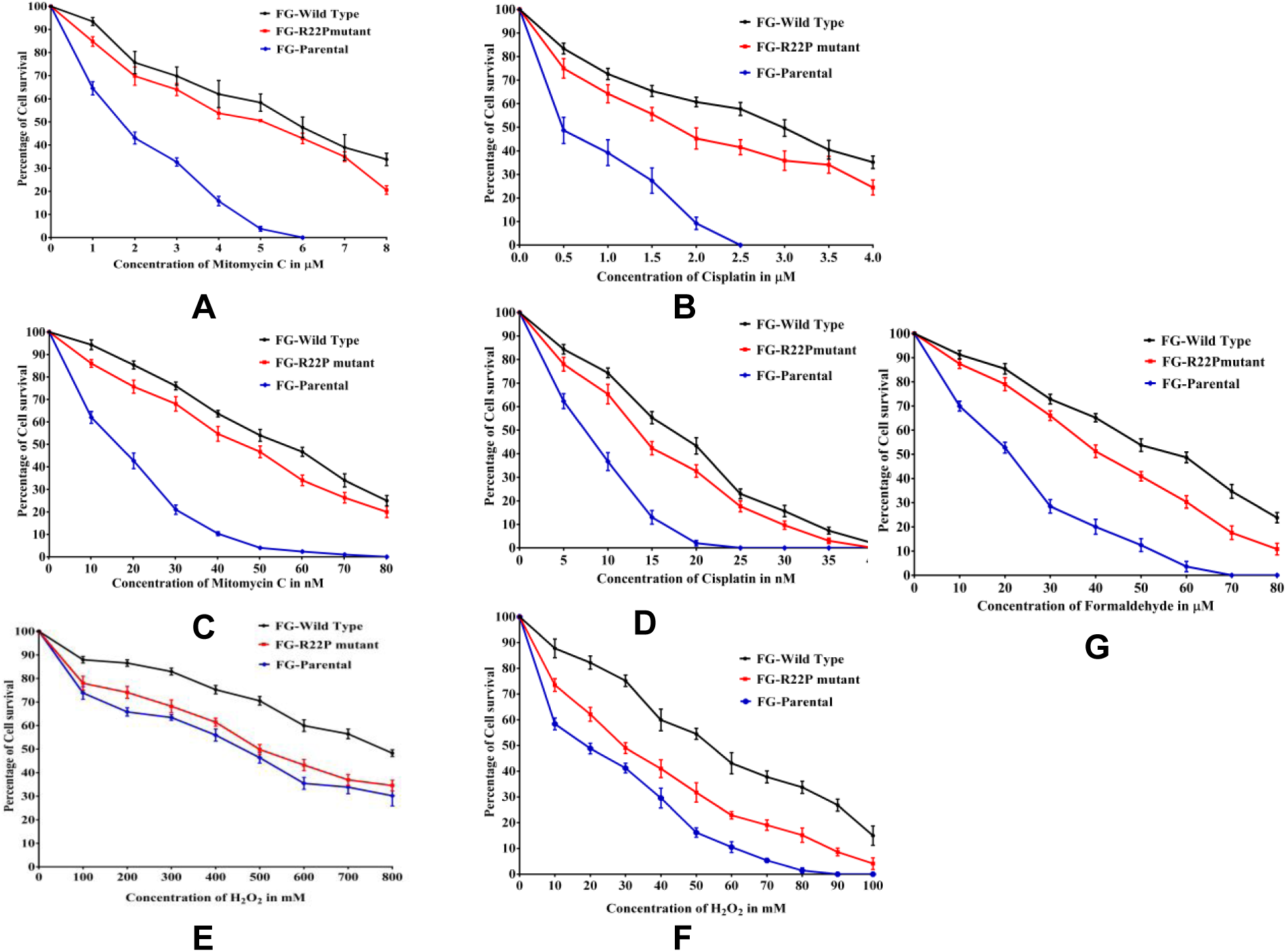
Drug sensitivity studies of FANCG corrected (black), FANCR22P (red) and FANCG parental cells (blue). Cells were treated with increasing concentration of drug (MMC and cisplatin) **(A & B)** for two days, and **(C & D)** five days, hydrogen peroxide (H_2_O_2_) **(E)** for two hrs and **(F)** twenty four hrs and **(G)** Formaldehyde for two hrs,. Cell survival was determined by MTT assay. Each value is the mean of three experiments.

In contrast, when the cells were treated with hydrogen peroxide for two hours and twenty-four hrs, like FANCG parental cells R22P cells showed sensitivity to oxidative stress (Fig.4E & F; supple Fig.S3 E & F). This result confirms our previous observation (Mukhopadhyay et al., 2006) that the role for the FANCG protein in mitochondria with respect to sensitivity to oxidative stress results from diminished peroxidase activity. In summary, it can be concluded that FANCG has dual roles: DNA damage repair in the nucleus and oxidative stress metabolism in mitochondria.

### Correlation between mitochondrial instability and genomic instability

The R22P mutant can repair the genomic DNA but fails to protect the mitochondria from oxidative stress. In spite of the nuclear DNA damage repair ability, the R22P patients are susceptible to getting cancer (D. et al., 2003). However, the question remains whether oxidative stress-mediated mitochondrial dysfunction influences the genomic DNA damage or not. Mitochondria of the R22P patients are under constant (endogenous) oxidative stress since birth and oxidative stress increases with age. Their genomic DNA is also attacked by several exogenous and endogenous ICL agents. Thus, an experiment on cell lines was set up to determine the extent of DDR in cells expressing the R22P mutant protein. The R22P cells, FANCG corrected cells, and FANCG parental cells were treated with mild oxidative stress (10µM of H_2_O_2_) for fourteen (14) hrs continuously, and at an interval of every two hrs, the cells were treated with a low dose of MMC (100nM) for thirty min. Then the cells were stained with JC-1 dye to determine the loss of mitochondrial membrane potential (ΔΨ), and γ-H2AX foci formation was determined in order to analyze the nuclear DNA damage (Fig. 5A). The percentage of depolarized (green and yellow) mitochondria and the number of nuclear foci at each time point were calculated in all three types of cells. From these results, we determined the percentage of functional mitochondria and the number of nuclear foci of the cells (8-10 fields and each field contains10-12 cells) as represented in graphical form (Fig. 5B). In FANCG corrected cells, the percentage of depolarized mitochondria is very low at initial time points (up to six hrs). Eight hrs onwards the percentage of depolarization started to increase, and by fourteen hrs approximately twenty percent of depolarized mitochondria are observed (Fig.5B). However, in both FANCG parental and R22P stable cells the percentage of depolarized mitochondria is very high at early time points (up to six hrs) compared to corrected cells. The percentage of depolarization is also increased with time in both cells. At fourteen hrs approximately 35 and 40 percent of depolarization mitochondria are observed in R22P and FANCG parental cells respectively (Fig. 5A&B). So, these experiments suggest that oxidative stress-mediated mitochondrial dysfunction is very high in FANCG parental and R22P cells compared to FANCG corrected cells.

**Fig. 5.**
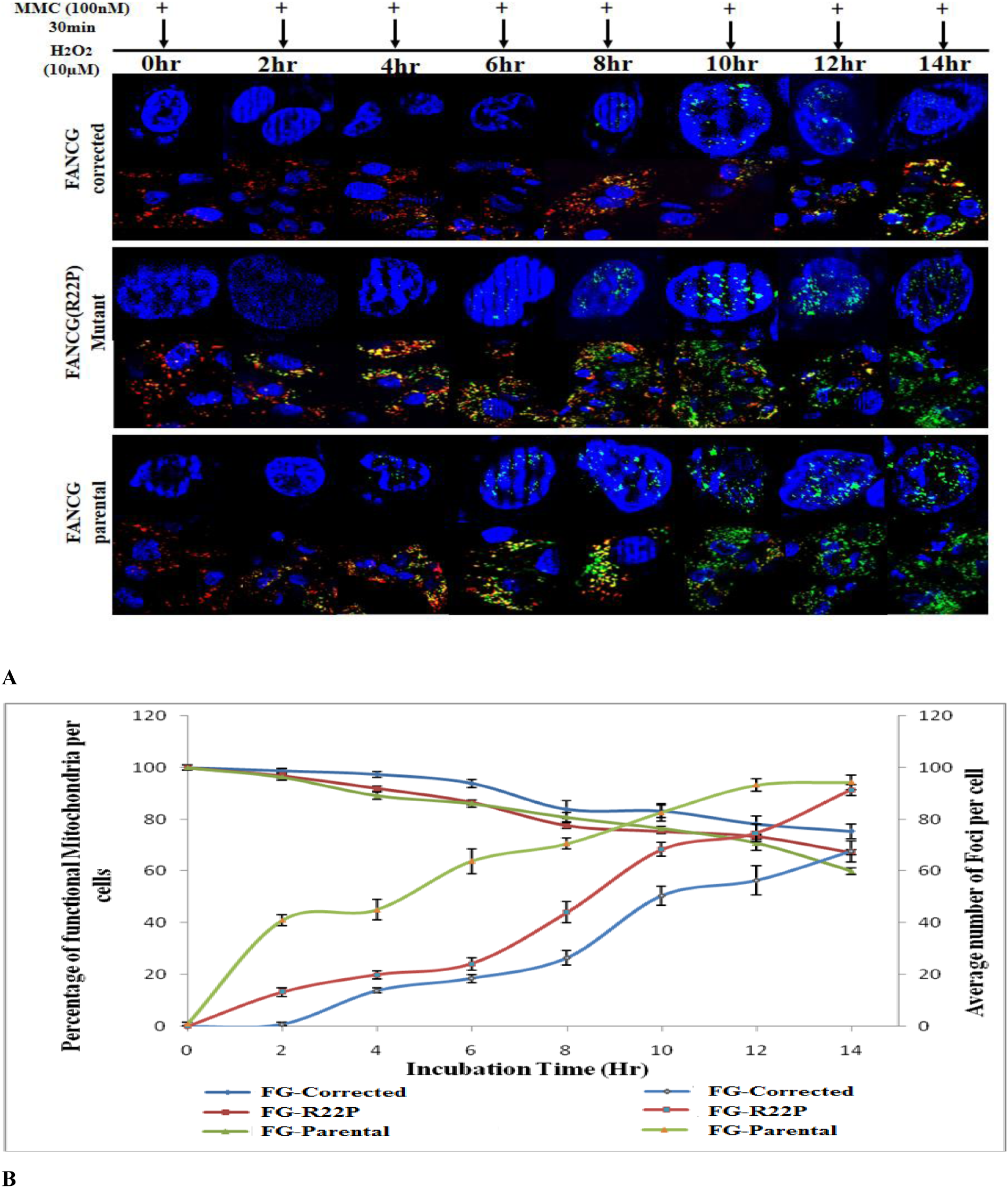
Mitochondrial depolarization and nuclear DNA damage in FG-corrected, FG-parental and FANCG-R22P cells. Cells were treated with H_2_O_2_ (10µM for fourteen hrs) and MMC (100nM for 30 min) at two hr-intervals. Nucleus was stained with γH2AX antibody and mitochondria were stained with JC-1. **(A)** The green dots in the nucleus represent the γH2AX foci. Red colour represents the normal mitochondria, green colour represents the depolarized mitochondria and yellow represents the intermediate values. Arrows represent the time of MMC treatment.**(B)** The graph represents the percentage of functional mitochondria and average number of foci in each type of cell. The values are the mean of multiple counts (more than three).

Similarly in FANCG corrected cells, the number of γ-H2AX foci is very low at early time points, and then the foci number increased with time. However, compared to the other two cells, the number of foci is low in FANCG corrected cells. At fourteen hrs the intensities of the foci are diminished, which suggests improved repair in the cells at that time point which was not observed either in FANCG parental or R22P cells (Fig. 5A). In FANCG parental cells, the foci number is very high at the initial time point of two hrs, and it continued to be high as compared to both FANCG corrected and R22P cells (Fig. 5A & 5B). The FANCG protein is absent in parental cells and is unable to protect either the nuclear DNA or the mitochondria. Whereas, in R22P cells, the number of foci is less at the initial time points of two to twelve hrs as compared to parental cells. After that, the number of foci is almost equal in both cell lines at a later stage (fourteen hrs; Fig 5B). As the foci number in the nucleus of R22P cell is lower than the parental cell at early times of treatment (2-12 hrs), this suggests that R22P cells can repair the DNA at early stages. After that percentage of depolarized mitochondria increased, crosses an apparent threshold, the R22P cells failed to repair DNA. Thus, at later times after exposure (14hrs) the number of foci is almost equal in both FANCG parental and R22P cells. All these observations clearly suggest that mitochondrial dysfunction influences the nuclear DDR.

### Mitochondrial instability causes defective FANCJ in R22P cells

Nuclear genomic instability can be the result of various types of mitochondrial dysfunction(Tokarz and Blasiak, 2014). One of the most important is the loss of mitochondrial membrane potential (ΔΨ) which inhibits the production of iron-sulfur prosthetic groups and impairs the assembly of Fe–S proteins (Kaniak-Golik and Skoneczna, 2015). Mitochondrial dysfunction inhibits the production of the iron-sulfur cluster (ISC) containing proteins, which are essential for maintaining the nuclear genome stability(Kaniak-Golik and Skoneczna, 2015; Lill et al., 2012; Richardson et al., 2010). To our surprise, we found that FA subtype J (FANCJ), an ISC containing helicase is essential for ICL repair. An attempt was made to see the transcriptional down-regulation of several of ISC-containing proteins involved in DNA damage repair along with FANCJ by real-time PCR. In our initial experiment the transcriptional down regulation was not observed in the cells (FANCG corrected, FANCG R22P and FANCG parental) treated with H_2_O_2_ (10µM) for fourteen hrs and followed by thirty min of MMC treatment for every two hrs (data not shown). Several human pathological mutants are identified in the Fe-S domain of FANCJ. The loss of iron-binding in the mutant protein resulted in the loss of helicase activity, suggesting the importance of iron in maintaining the structure and function of FANCJ (Wu et al., 2010). We wanted to compare the status of the FANCJ protein in terms of iron-binding and helicase activity in all three sets of cells at each time point of the experiment (10µM of H_2_O_2_ for fourteen hrs and followed by thirty min treatment with100 nM of MMC after every two hrs). Cells were treated with medium containing labeled iron (^55^Fe). After MMC treatment cells were lysed at every time point. An equal amount of protein was used for IP with FANCJ antibody and protein A/G agarose beads. The quantitation of ^55^Fe was used to estimate the amount of iron in FANCJ(Pierik et al., 2009). The amount of iron present in FANCJ of each cell type at zero time was considered as hundred and was compared with the amount of iron present in the same cells at other time points (relative percentage). The continuous reduction of iron of FANCJ with time was observed only in both the R22P and FANCG parental cells (Fig.6A). Almost a fifty percent loss of iron of the FANCJ protein was observed in these two cells at later stages (8,10,12 and 14hrs) (Fig.6A) compared to FANCG corrected cells. Whereas, the percentage of iron of FANCJ was unaltered or was increased in FANCG corrected cells. Western blot analysis confirmed the amount of FANCJ protein present in each IP(Fig.6B). From these observations, it can be concluded that loss of mitochondrial membrane potential (ΔΨ) causes iron depression of the FANCJ protein only in R22P stable and FANCG parental cells. FANCJ is dependent on mitochondria for Fe-S cluster domain (Rudolf et al., 2006). As a control iron binding of ferritin is not reduced in these experiments (Fig. S4). Ferritin is independent of mitochondria for the iron source (Mackenzie et al., 2008). Thus, the *in vivo* iron uptake experiment suggests that the high percentage of depolarized mitochondria causes the iron deficiency of FANCJ in both R22P and FANCG parental cells which potentially may affect the helicase activity.

**Fig. 6.**
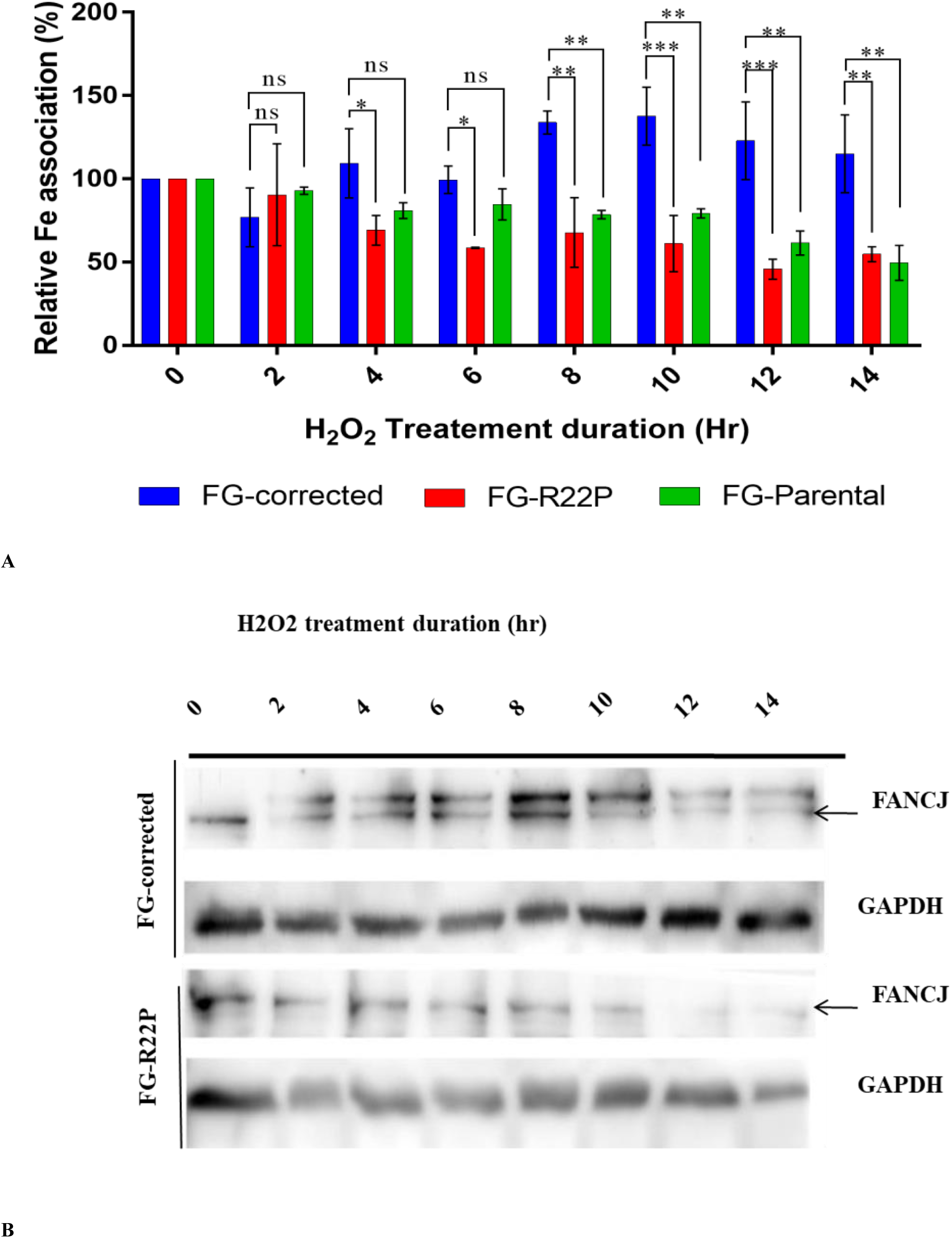
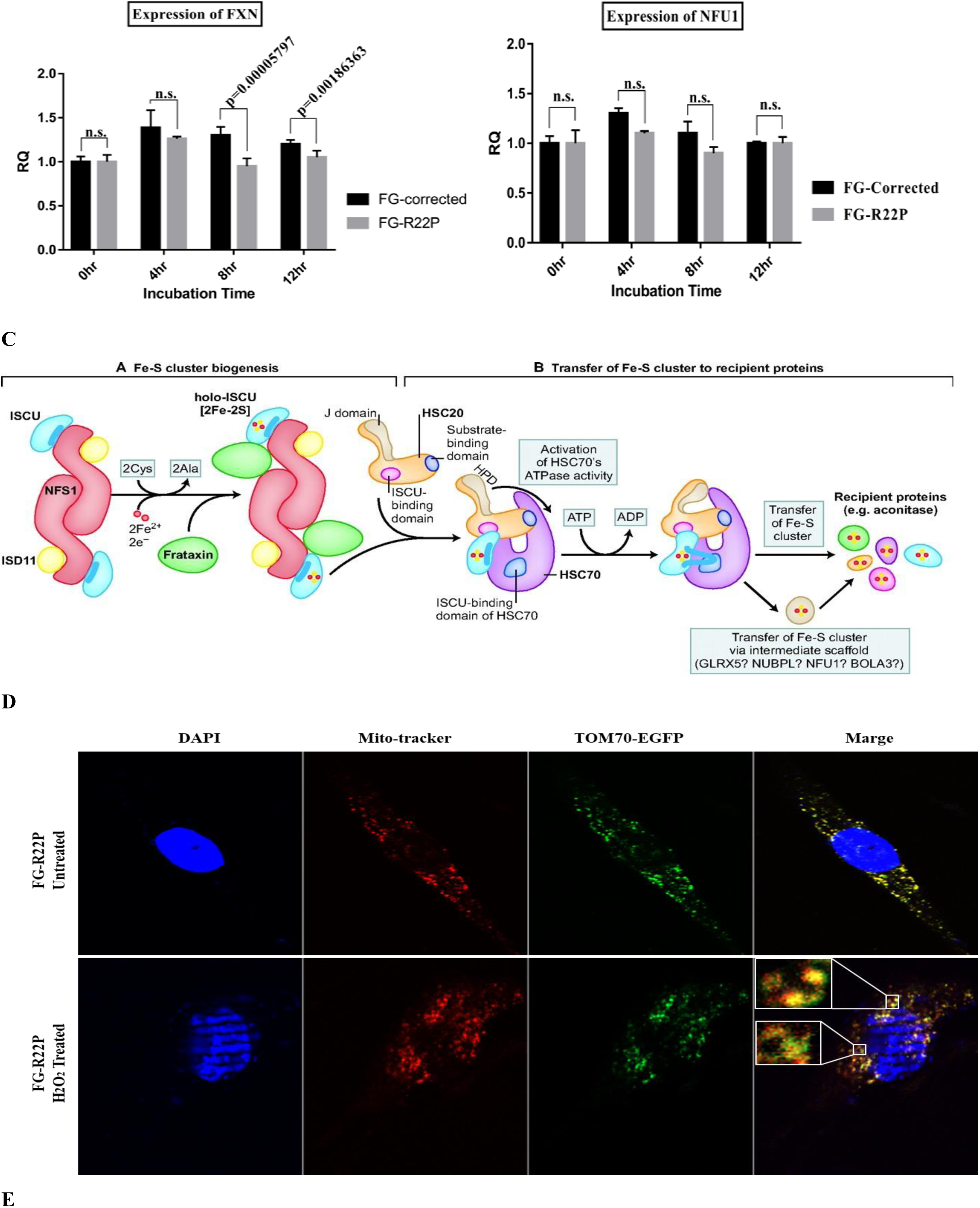
Quantification of the amount of iron present in FANCJ protein of FG-corrected, FG-parental and FG-R22P cells. **(A)** Amount of ^55^Fe at 0 hr was considered as hundred and relative amount of ^55^Fe was calculated at each time point. Each result is the mean of minimum three experiments.(ns = non-significant, *=0.01<p<0.05, **= 0.0001<p<0.0009, ***= p≤0.0001 for α=0.05) **(B)** Western blot of the cell lysates with FANCJ and GAPDH antibody. (C) mRNA expression of FXN and NFU1at different time points. (D) Diagram of Fe-S clustur biogenesis and transfer to recipient proteins () (E) Co-localization of TOM70-GFP and mitotracker in HeLa cell.

Fe-S cluster metabolism occurs in mitochondria in two major steps: (i) Fe-S cluster synthesis and (ii) transfer of Fe-S cluster to recipient protein (Fig6D). A complex of proteins is involved in each step(Lill and Mühlenhoff, 2008). The Fe-S cluster biogenesis proteins are mainly nuclear protein, and they migrate to mitochondria. The cause of iron deficiency of FANCJ protein in R22P and FANCG parental cells is either the difficulty of migration of the nuclear protein into mitochondria due to alteration of mitochondrial membrane potential or the downregulation of Fe-S cluster proteins due to mitochondrial stress. Cytochrome C is nuclear gene but present in the mitochondrial matrix. Tom70 is nuclear gene but present in the mitochondrial outer membrane. The localization of Cytochrome C may alter due to loss of mitochondrial membrane potential. But, the localization of Tom70 should not be hampered in spite of mitochondrial membrane potential loss (Veatch et al., 2009). To help elucidate this mechanism we have transiently expressed human Tom70-tagged with GFP and human Cytochrome C-tagged with RFP into the R22P cells. Cells without treatment showed complete co-localization of both Tom70 and Cytochrome C. But the cells treated with H_2_O_2_ did not show complete co-localization of Tom70 with Cytochrome C (Fig.6E). Thus, these results suggest that Cytochrome C did not migrate into the matrix due to the loss of mitochondrial membrane potential. We have also studied the transcriptional expression of the Fe-S cluster genes in the FANCG corrected and R22P cells. The cells were treated with 10µM of H_2_O_2_for twelve hrs and followed by thirty min treatment with100nM of MMC after every two hrs. The expression of the genes was compared between these two cells by Real Time PCR (Fig.6C). The expression of Frataxin (Fxn) was significantly decreased in R22P cells as compared to FANCG corrected cells. The lower expression of FXN was observed from 8hrs of treatment, consistent with the result shown in Fig.6A. FXN is essential for Fe-S cluster biogenesis. However, the expression of NFU1 is not altered (Fig6C). Nfu1 is responsible for transportation of the Fe-S cluster to recipient proteins. Thus these results suggest that the transcription of the FXN is significantly reduced in R22P cells as compared to FANCG corrected cells, because of the greater number of dysfunction mitochondria in R22P cells. However, decreased mitochondrial migration of the ISC proteins due to loss of mitochondrial membrane potential also cannot be ruled out.

## Discussion

### Unique Mitochondrial Localization Signal of human FANCG

We have used different in silico tools to identify the mitochondrial localization signal of human FANCG. These tools predict two things; (i) whether the protein contains any mitochondrial localization signal or (ii) mitochondrial localization of the protein. These analyses strongly predicted the N-terminal thirty amino acids of human FANCG as a mitochondrial localization signal (MLS) and correlated with the mitochondrial existence of the human FANCG. FANCG sequences from other species have been analyzed, and none of them are found to carry the MLS/mTP either at N-terminal or C-terminal except human. Thus, one hypothesis to explain this discrepancy is that the MLS region has evolved later in humans and as a result, human FANCG has acquired the ability of regulates mitochronrial function in addition to the nuclear DNA damage repair.

Interestingly, the FANCG knockout(KO) mice do not exhibit any severe phenotype. FANCG cells derived from KO mice are only mildly sensitive to ICL agents, but not sensitive to oxidative stress (Parmar et al., 2009; Pulliam-Leath et al., 2010; Yang et al., 2001). However, expression studies of FANCG in other species will be required to help explain this result.

We have identified several mutations in the MLS region of human FANCG from the LOVD, and COSMIC (catalogue of somatic mutation in cancer) database and their mitochondrial localization has been studied (unpublished results). In this report, we are describing one pathogenic mutation where the 22^nd^ Arginine has been replaced by Proline (FANCGR22P). GFP expression of this fusion construct suggests its inability to migrate into mitochondria. The predicted structure suggests as expected by many studies of the effects of proline insertions on alpha-helical structures that the helix is broken due to replacement of arginine by proline at the N-terminal of FANCG. Several studies suggested the importance of arginine for mitochondrial localization of proteins (Neve and Ingelman-Sundberg, 2001). A most interesting feature of this mutant protein combines its inability to migrate to mitochondria with its ability to translocate to the nucleus. These phenotypes were confirmed by drug sensitivity experiments. R22P stable cells are resistance to ICL drugs like FANCG-corrected cells and sensitive to oxidative stress like FANCG parental cells. FANCD2 monoubiquitination in R22P cells also suggests the ability of the mutant protein to form the FA complex. Thus, the phenotype of the R22P pathogenic mutation resolves the long-lasting debate of FA protein’s role in mitochondria. An open question can be raised about the implication(s) of these results in the clinical diagnosis of FA patients. Some patients are diagnosed as FA by phenotypic features, though the drug sensitivity test of their cells suggests negative (chromosome breakage test). In that case, drug sensitivity tests should be performed in the presence of mild oxidative stress (Suppl Fig.5A, 5B)

### Mitochondrial dysfunction causes defective FANCJ: Mitochondrial instability leads to genomic instability

The inability of the cell to repair DNA damage may result in cancer. In this study, we have found that despite the genomic DNA repair ability, the R22P patients are also affected by cancer (LOVD database). R22P cells are highly sensitive to oxidative stress, and loss of mitochondrial membrane potential is observed due to oxidative stress (Mukhopadhyay et al., 2006). From these two observations, we suggest that there is a correlation of the mitochondrial instability with genomic instability. Many studies suggest that mitochondrial DNA mutation and loss of mitochondrial membrane potential may cause cancer (Tokarz and Blasiak, 2014). One proposed mechanism is that the reactive oxygen species (ROS) produced due to mitochondrial dysfunction may destabilize the cellular macromolecules, including the damage of genomic DNA(Nunnari and Suomalainen, 2012). So far, an association of oxidative stress with inter-strand cross-linking (ICL) damage is not known. In order to elucidate this association, we performed an experiment with R22P cells (Fig.5). The results suggest that R22P cells can repair the ICL damage as long as there is a certain level of functional mitochondria in the cell. When this percentage is reduced, R22P cells fail to protect their genomic DNA from ICL damage. Fe-S containing proteins are essential for their role in various cellular functions such as catalysis, DNA synthesis, and DNA repair (Netz et al., 2014). Fe-S proteins depend on mitochondria for their Fe-S domain because the iron-sulfur cluster (ISC) synthesis is one of the major functions of mitochondria (Lill and Mühlenhoff, 2008). Several reports suggested that assembly of all ISC-containing proteins requires intact mitochondria(Biederbick et al., 2006). Even the loss of mitochondrial DNA or loss of mitochondrial membrane potential impairs the ISC biogenesis(Kispal et al., 1999). Recently, Daniel Gottschiling’s group has shown in a yeast system that loss of mitochondrial DNA causes a defect in mitochondrial iron metabolism(Veatch et al., 2009). But this study is the first report of defective Fe-S containing protein FANCJ due to oxidative stress-mediated mitochondrial dysfunction. In vivo studies in R22P cells suggest that significant deficiency of iron in FANCJ helicase occurs due to loss of mitochondrial membrane potential (Fig.6A). Several studies suggest that deficiency of iron of the Fe-S containing protein may result in the loss of helicase activity(Wu et al., 2010), but we were unable to test this directly. Moreover, in our studies, we only have studied FANCJ, but other Fe-S containing cellular proteins involved in DNA damage repair also might have been affected (Netz et al., 2014). Altogether, our results strongly suggest that in normal cells FANCG protects the mitochondria from oxidative stress. As a result, mitochondria maintain the ISC biosynthesis and provide Fe-S cluster for maintaining the Fe-S domain of active FANCJ helicase, which is required for nuclear DNA damage repair (Fig7). In R22P cells, ISC biosynthesis is either low or impaired due to mitochondrial dysfunction. We have identified the down regulation of FXN in R22P cells as compared to FANCG corrected cells under stress condition. FXN is an important protein involved in ISC biogenesis, and the defect in ISC biosynthesis leads to various human diseases. However, how the mitochondrial stress regulates the transcription of Fxn is not known. One possibility is that via SP1, a ubiquitous transcription factor present in the promoter region of the human Fxn gene(Li et al., 2010). The sumoylation of SP1 under oxidative stress and the subsequent lack of DNA binding has been reported(Wang et al., 2008).

**Figure 7.**
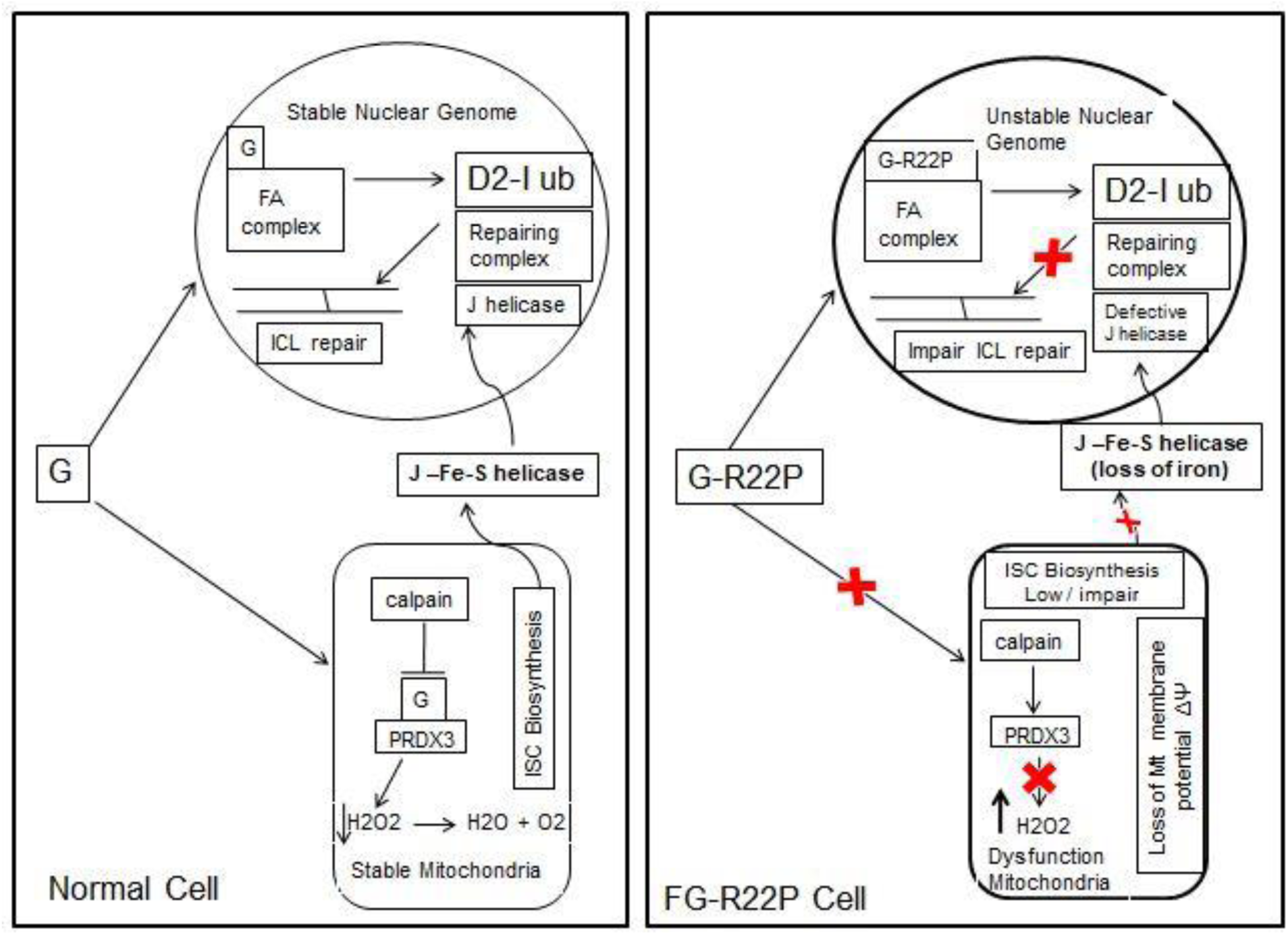
Model to Explain the mitochondrial instability leads to genomic instability. **(A)** In normal cell FANCG prevents PRDX3 from calpain cleavage, and maintains mitochondrial stability by reducing oxidative stress. Stable mitochondria maintain the helicase activity of FANCJ by providing ISC domain. **(B)** In FG-R22P cell FANCG fails to migrate to mitochondria and PRDX3 is cleaved by calpain. Mitochondrial membrane potential (ΔΨ) is lost due to elevated oxidative stress and ISC biosynthesis is reduced. FANCJ lost its helicase activity due to insufficient iron in its Fe-S domain

The difficulty in mitochondrial import of nuclear proteins involved in ISC biosynthesis provides another possibility. As a result, FANCJ will be depleted in iron required for Fe-S domain. So, the defective FANCJ is unable to repair the nuclear DNA (Fig.7). We found that there is certain percentage of defective mitochondria in the cells which do not affect the overall ISC biosynthesis in the cell. However, when this number decreases to a critical threshold, then the ISC containing proteins will undergo an iron crisis. Further studies with R22P cells are required to identify the threshold percentage of dysfunctional mitochondria. Our studies with certain FA mutations help confirm the relevance of non-respiratory function of mitochondria in disease progression. This is not unique, but a common phenomenon, the known consequence of cellular oxidative stress.

### Experimental Design

#### In-silico tools used

1. TargetP1.1(http://www.cbs.dtu.dk/services/TargetP/)
2. iPSORT server (http://ipsort.hgc.jp/)
3. MitoProt(https://ihg.gsf.de/ihg/mitoprot.html)
4. PredotarMito(https://urgi.versailles.inra.fr/predotar/predotar.html)
5. TPpred2.0(http://tppred2.biocomp.unibo.it/tppred2/default/help)
6. RSLpred(http://www.imtech.res.in/raghava/rslpred/)
7. iLocAnimal(http://www.jci-bioinfo.cn/iLoc-Animal)
8. MultiLoc/TargetLoc(https://abi.inf.uni-tuebingen.de/Services/MultiLoc)

#### Database used

1. LOVD database(http://databases.lovd.nl/shared/variants/FANCG/unique)
2. COSMIC database (http://cancer.sanger.ac.uk/cosmic/gene/analysis)

#### Protein structure prediction

1. Secondary structure: An advanced version of PSSP server is used for prediction of protein secondary structure by using nearest neighbor and neural network approach.
2. Tertiary structure: the Tertiary structure is predicted by I-TASSER(http://zhanglab.ccmb.med.umich.edu/I-TASSER/). Structural templates of the proteins are first identified from the PDB by multiple threading approach LOMETS. Full-length atomic models were constructed by iterative template fragment assembly simulations. The function insights of the target proteins were finally derived by threading the 3D models through protein function database BioLiP.

#### Cell lines

The cell lines HeLa and HEK293 were obtained from ATCC and maintained in Dulbecco’s modified Eagle medium (DMEM) supplemented with 10%(v/v) fetal bovine serum(FBS) and 1x penicillin/streptomycin (Himedia). FANCG corrected (FG^+/+^) and FANCG parental (FG^-/-^) fibroblast cells were generous gift from Dr.AgataSmogorzewska (The Rockefeller University, New York, USA) while FANCG mutant(R22P) fibroblast cells were prepared in our laboratory and are maintained in Dulbecco’s Modified Eagle medium (DMEM) supplemented with 15% (v/v) fetal bovine serum (FBS) and 1x penicillin/streptomycin (Himedia).

#### Antibodies

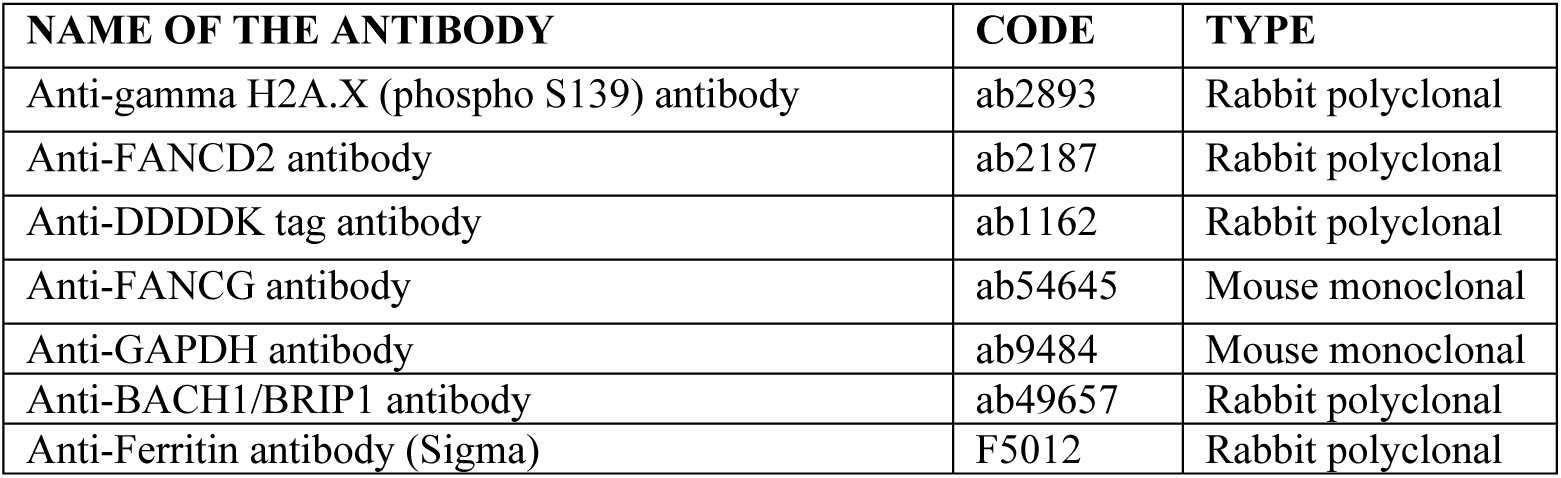

**Antibodies:**
| NAME OF THE ANTIBODY | CODE | TYPE |
| --- | --- | --- |
| Anti-gamma H2A.X (phospho S139) antibody | ab2893 | Rabbit polyclonal |
| Anti-FANCD2 antibody | ab2187 | Rabbit polyclonal |
| Anti-DDDDK tag antibody | ab1162 | Rabbit polyclonal |
| Anti-FANCG antibody | ab54645 | Mouse monoclonal |
| Anti-GAPDH antibody | ab9484 | Mouse monoclonal |
| Anti-BACH1/BRIP1 antibody | ab49657 | Rabbit polyclonal |
| Anti-Ferritin antibody (Sigma) | F5012 | Rabbit polyclonal |

#### Constructs

pcDNA3-EGFP (#13031), pLJM1-EGFP (#19319), pCMV-VSV-G (#8454) and pSPAX2 (#12260) was obtained from Addgene. The construct pDsRed2-Mito encoding a fusion of *Discosomasp* red fluorescent protein (DsRed2) and a mitochondrial targeting sequence of human cytochrome c oxidase subunit VIII (Mito) was purchased from Clontech Laboratories. Full-length FANCG-wt cDNA was initially cloned into TA vector pTZ57R/T (Thermo-scientific) that was further utilized as a template for the full length and N-terminal deletion constructs of FANCG. Full length and N terminal deletion (up to 30 amino acid) constructs of FANCG were subcloned into the KpnI and EcoRI site of pCDNA3-EGFP to encode C-terminal EGFP tagged FANCG proteins. FANCG mutants were constructed by conventional PCR method, using full-length FANCG-wt-pcDNA3-EGFP as a template followed by DpnI treatment. FANCG mutant R22P (Arginine to proline at the 22^nd^ position from N-Terminal) was further sub cloned into the EcoRI and SpeI site of lentiviral vector pLJM1-EGFP. Full length TOM70-wt was cloned into KpnI and EcoRI site of pcDNA3-EGFP to encode C-terminal EGFP tagged TOM70 proteins. All the primers utilized are given below.

#### Primers

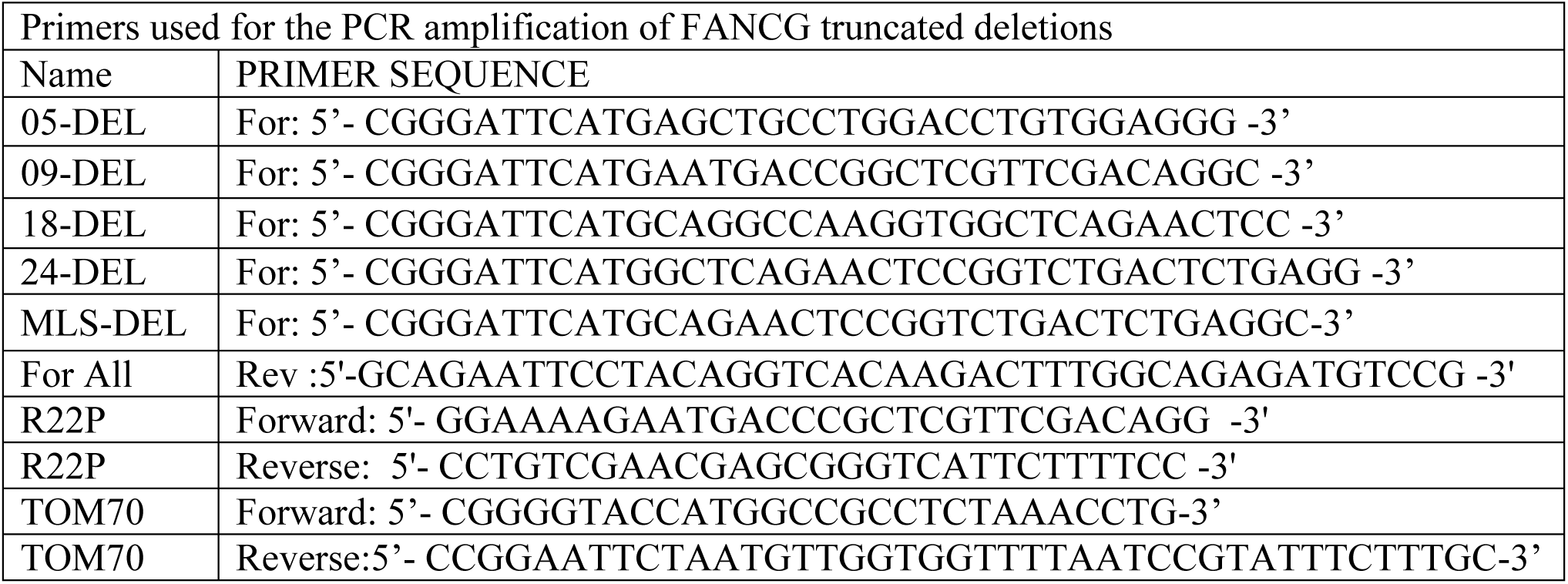

### Immunofluorescent Microscopy

Cells were grown onto Poly-L-Lysine coated coverslips in 60mm dishes and were transfected with the indicated constructs using Lipofectamine(Fermentus) for 48hrs. Cells were incubated in blocking buffer (5% nonfat milk in 1XPBS/ 5% FBS in 1XPBS) for one hr followed by one hr incubation with primary antibody at room temperature. After that cell was washed with 1X PBS followed by one hr incubation with respective secondary antibody tagged with either FITC or Texas Red. The cells were fixed with either with 4% paraformaldehyde (Himedia) for 10 minutes or in ice-cold methanol solution for 5 minutes. 0.2% TritonX-100 can be treated for two minutes for permeabilization of the antibody into cells. Then the coverslips were air dried and mounted with mounting medium with DAPI (Vectashield) on a glass slide by inverting the coverslips upside down. The mounted cells kept in dark for 15 minutes and fixed the coverslip with transparent nail polish. Imaging was performed on a fluorescence microscope (Axio observer.Z1, Carl Zeiss Micro-Imaging, Germany) attached with Axiocam HRM CCD camera and Apotome.2. Axiovision software (Zenpro2012) and Adobe photoshop7.0 software were used for deconvolution imaging and image analysis.

### FANCG R22P Mutant Stable Cell Line development

#### Development of Virus Particle in HEK293 Cells

FANCG R22P initially was cloned into pCDNA3-EGFP. It was PCR cloned into the TA vector pTZ57R/T (Thermo Scientific) for creating compatible enzyme site for Lenti vector. Now the R22P construct was digested with EcoR1 and BamH1and cloned into the viral packing vector pLJM1-EGFP (Addgene). The viral particle protein containing vectors pCMV-VSV-G, psPAX2, and FG R22P-pLJM1-EGFP constructs were transfected (1:1:3) into HEK293 cells by Turbofect (Fermentas). The cells were grown for forty-eight (48) hrs for development of the virus particle.

#### Integration of R22P into the genome of FANCG (-/-) parental cells

The cell culture medium containing virus particles were collected into a 15 ml sterile centrifuge tube and centrifuged 14,000rpm for 30 minutes at 4°C to remove the cellular debris and stored at 4°C. Fresh media was added to each HEK293 cell monolayer and incubated for another 12 hrs. This media was collected and centrifuged 14,000 rpm for 30 minutes at 4°C and mixed with the earlier supernatant. Total media was filtered by 0.22-μm syringe filter unit (Millipore) and was centrifuged again for 60 min at 14,000 rpm, at 4°C. Gently the supernatant was removed by pipette and fresh media was added to the tube containing precipitate at the bottom and again the same step was repeated to concentrate the lentiviral particles. Ultimately the supernatant containing virus was added to the flask of sixty (60) percent confluent FANCG parental cells and incubated the cells for forty-eight hrs. Two days after infection, the cells were checked for GFP fluorescence and puromycin resistance cells were developed by adding increasing concentrations of puromycin (2-5μg/ml) in the media. The puromycin resistant cells were subcultured several times and preserved in freezing media at −80°C for further usage. The stable cells were confirmed by Western blot with FANCG specific antibody.

### Cell Survival Assay

#### Cell Viability Test by Trypan blue dye exclusion

An equal number of each cell (FANCG corrected, FANCG parental and R22p FANCG stable) were seeded in 8 well tissue culture plates with respective blank and controls for twenty-four hrs. The DNA damaging agents (MMC, Cisplatin, H_2_O_2_ and Formaldehyde, their concentrations were mentioned as below) were added to each well and incubated for 24 hrs. 0.1 mL of 0.4% solution of Trypan blue in buffered isotonic salt solution, pH 7.2 to 7.3 (i.e., phosphate-buffered saline) (Himedia) was added to 1 mL of cells. These cells were loaded on a hemocytometer and the cells were counted in each well for the number of blue staining cells and the number of total cells. Cell viability nearly 95% maintained for healthy log-phase cultures. In order to calculate the number of viable cells per mL of culture the formula below was used.

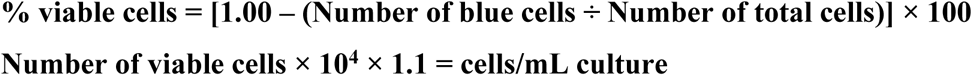

The viable cells were used for the preparation of the graph at each concentration of the drug. Triplicate experiments were performed for each concentration.

#### Cell Cytotoxicity Test by MTT assay

An equal number of cells (FANCG corrected, FANCG parental and R22P FANCG stable) were seeded into 96 well tissue culture plates with respective blank and control and incubated for twenty-four hrs. The DNA damaging agents (MMC, Cisplatin, H_2_O_2_ and Formaldehyde, their concentrations were mentioned as below) were added to each well and mix by gently rocking several times and incubated for 48 hrs. 20μl of MTT reagent (Thiazolyl Blue Tetrazolium Bromide) at 5mg/ml concentration in sterile PBS (Himedia) was added to each well and mixed by gentle rocking and incubated for 1 hour. The media was removed without disturbing cells and purple precipitate. 200μl of DMSO was added for solubilization of the purple precipitate formazan. The plate was shaken at 150rpm for 10 minutes for equal mixing the formazan into the solvent. The optical density of the solution was observed at 570nm on the ELIZA plate reader.

#### GammaH2Ax foci Assay

FANCG corrected(FG^+/+^), FANCG parental (FG^-/-^) and FANCG mutant (R22P) fibroblast cells were plated onto Poly-L-Lysine coated coverslips in 60mm dishes in DMEM with 15%(v/v) FBS and 1X penicillin/streptomycin solution and were kept in 5% CO2 incubator followed by serum starvation upon achieving nearly 60 to 70 percent confluency. These cells were treated with H_2_O_2_ (10µM) in serum-free DMEM for continuous fourteen hrs followed by MMC (100nM) treatment for 30min in every 2hr interval. Treated cells were incubated with rabbit polyclonal anti-phospho-gamma H2AX antibody (ab11174). Goat-anti-rabbit IgG daylight secondary antibody (Thermo) was used for 1 hour followed by DAPI staining and foci were calculated under a microscope (Axio observer.Z1, Carl Zeiss MicroImaging, Germany) equipped with Axiocam HRM CCD camera and Apotome.2. Axiovisionsoftware (Zenpro2012).

#### Determination of mitochondrial membrane potential loss with JC dye

Same FANCG fibroblast cells which were analyzed for the GammaH2Ax foci have also analysed for the mitochondrial membrane potential. At every two hrs after treatment with MMC, the cells were stained with 1XJC1 dye solublised in DMSO for 10 minutes. Then the cover slips were air dried and mounted with Vectashield mounting medium. The mounted cells kept in dark for 15 minutes and fixed the cover slip with transparent nail polish. These cells were observed under Zeiss microscope with DAPI, FITC and PI filters for mitochondrial membrane potential change and deconvoluted images captured with Fluorescence microscope (Zeiss Axio observer.Z1) fitted with Axiocam observer camera. Red color represents the stable mitochondria. Green represents the loss of mitochondrial potentiality. Yellow is the in-between status of red and green.

### In vivo iron uptake assay of FANCJ(Pierik et al., 2009)

Fibroblast cells (FANCG corrected, FANCG parental and R22P stable cell) were grown at 37°C with 5% CO2in Iscove’s Modified Dulbecco’s Medium (IMDM, Sigma) supplemented with 15% (v/v) fetal bovine serum(FBS, Gibco), and 1X penicillin/streptomycin solutions(Gibco) for 24hr. Followed by second incubation for 2hr in IMDM containing 15% (v/v) FBS,1X pen-strep and 10µCi ^55^Fe (BARC, India).Following the serum starvation in IMDM containing 10µCi ^55^Fe for 2hr, these cells were treated with H_2_O_2_(10µM) in serum-free IMDM for continuous fourteen hrs followed by MMC (100nM) treatment for 30min in every 2hr interval. Cells were collected, washed three times with 1X PBS and were lysed in IP lysis buffer (25mM HEPES, 100mM NaCl, 1mM EDTA, 10% (v/v) glycerol, 1%(v/v) NP-40) supplemented with 1mM PMSF (phenylmethylsulfonyl fluoride), 10mM DTT (dithiothreitol), 1mM sodium orthovanadate,10ng/mL leupeptin, 1ng/mL aprotinin. Approx 700µg cell lysate was incubated with 4-5µl of IP grade polyclonal FANCJ antibody (Abcam) and ferritin antibody (Sigma) for 1hr at 4°C. Ferritin was taken as control for iron uptake by cells. 30 µl of protein A/G plus agarose beads (Biobharti, India) was added and incubated at 4°C for overnight under constant shaking.Beads were washed three to four times with 1X IP lysis buffer. Washed beads were boiling in 10%(w/v) SDS solution and were mixed with scintillation oil. DPM(disintegration per minute) of ^55^Fe were count in a liquidscintillation counter.

### DNA substrate

Standard desalted oligonucleotides were purchased from IDT and were used for the preparation of DNA substrates. The forked-duplex DNA substrate was prepared from the DC26 and TSTEM25 oligonucleotides as described by(Wu et al., 2010).

### FANCJ Helicase Assay

Fibroblast cells (FG^+/+^, FG^-/-^ and R22P mutants) were grown at 37°C with 5% CO_2_ in Dulbecco’s Modified Eagle Medium (DMEM, Gibco) supplemented with 15%(v/v) fetal bovine serum(FBS, Gibco), and 1X penicillin/streptomycin solutions(Gibco) for 24hr followed by serum starvation in DMEM for 2hr. These cells were treated with H_2_O_2_(10µM) in serum-free DMEM for continuous fourteen hrs followed by MMC (100nM)treatment for 30min. In every 2hr interval, Cells were collected, washed three times with 1X PBS and were lysed in IPlysis buffer (25mM HEPES, 100mM NaCl, 1mM EDTA, 10%(v/v) glycerol, 1%(v/v) NP-40) supplemented with 1mM PMSF (phenylmethylsulfonyl fluoride), 10mM DTT (dithiothreitol), 1mM sodium orthovanadate,10ng/mL leupeptin, 1ng/mL aprotinin. Approx 700µg cell lysate was incubated with 4-5µl of IP grade polyclonal FANCJ antibody(Abcam) for 2hr at 4°C, followed by 3hr incubation with 30 µl of protein A/G plus agarose beads (Biobharti, India) at 4°C under constant shaking. Beads where then collected by centrifugation at 4°C and washed two times with 1X IP lysis buffer and two times with 1X Helicase buffer (40mM Tris-HCl (pH 7.4), 25mM KCl, 5mM MgCl_2_, 0.1mg/ml BSA,2% (v/v) Glycerol, 2mM DTT). Helicase reaction was initiated by incubating FANCJ bound washed A/G plus Agarose beads at 37°C for 30min with helicase reaction mixture containing helicase buffer, 2mM ATP and 0.5nM of DNA substrate.was then incubated with Helicase buffer, 0.5nM DNA substrate. Reaction were terminated using stop buffer (0.3% w/v SDS and 10mM EDTA). The reaction product was resolved on nondenaturing 11% (30:1 acrylamide-bisacrylamide) polyacrylamide gel followed by drying and then were subjected to autoradiography.

### RNA isolation,cDNA synthesis and Real-time PCR

Fibroblast cells (FG^+/+^ and R22P mutants) were grown at 37°C with 5% CO_2_ in Dulbecco’s Modified Eagle Medium (DMEM, Gibco) supplemented with 15%(v/v) fetal bovine serum(FBS, Gibco), and 1X penicillin/streptomycin solutions(Gibco) for 24hr followed by serum starvation in DMEM for 2hr. These cells were treated with H_2_O_2_ (10µM) in serum-free DMEM for continuous twelve hrs., followed by MMC (100nM) treatment for 30min. At every 4hr interval, Cells were collected by trypsinization. Total RNA was isolated from these harvested cells using Trizol (Ambion, life technology) and then was stored in −80°C until further use.

4µg of above-isolated total RNA was used to prepare cDNA using the Verso cDNA Synthesis Kit (Thermo scientific), following the manufacturer’s protocol. Prepared cDNA was then diluted five times and 2 ul of this diluted cDNA was used as template for Real-time PCR.

Real-time PCR was permformed on StepOnePlus Real time PCR system (Applied biosystems) using the syber green PCR master mixture (Applied biosystems, Thermo Fisher Scientific). The program was set as follow; Holding Stage; 95°C, 10min, cycling stage; 40 cycle, 95°C, 15sec, 57°C, 1min and 60°C, 1min, Melting curve stage; step and hold, 95°C, 15sec, 60°C, 1min, 95°C, 15min with ramping rate of +0.3°C. β-Actin was used as endogenous control. The Sequence of Primers used for Real-time is given below. Graph Pad Prism 7 was used to perform multiple t-test to evaluate the statistical significance, using the Two-stage linear step-up procedure of Benjamini, Krieger and Yekutieli, with desire FDR(Q) = 5% without assuming a consistent Standard deviation.

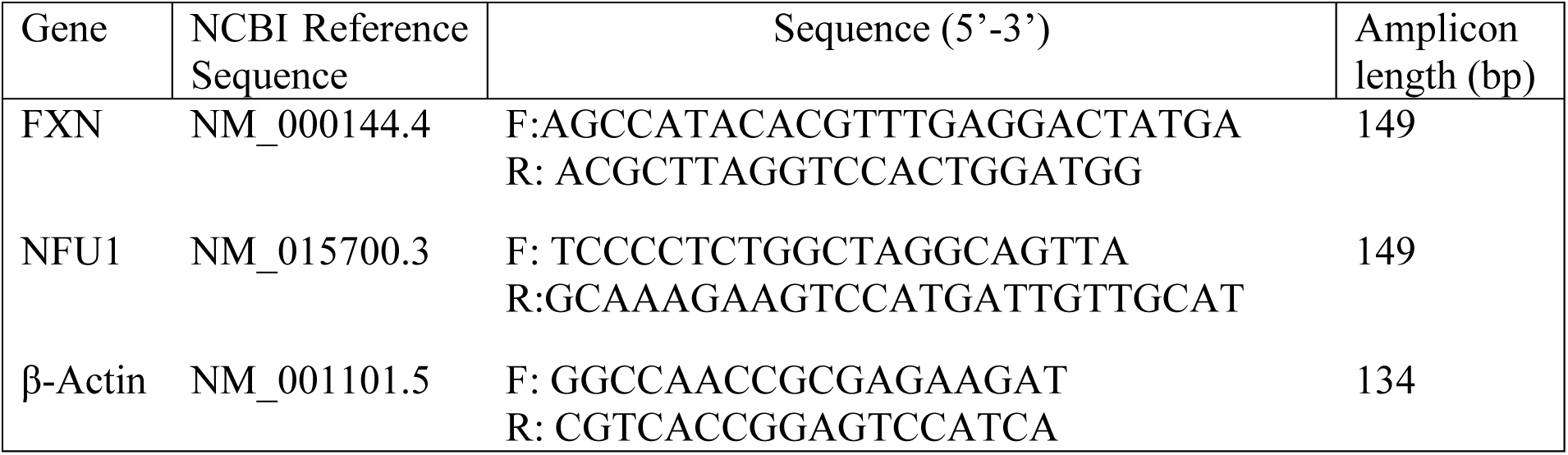

### Chromosome Preparation

Fibroblast cells (R22P mutants) were grown at 37°C with 5% CO_2_ in Dulbecco’s Modified Eagle Medium (DMEM, Gibco) supplemented with 15%(v/v) fetal bovine serum(FBS, Gibco), and 1X penicillin/streptomycin solutions(Gibco) for 24h, followed by serum starvation in DMEM for 2hr. These cells were treated with either MMC (100nM) or H_2_O_2_ (300µM) and MMC (100nM) in serum-free DMEM for continuous two hrs, Followed by colcimeid (200µg/ml) treatment for 1hr. These cells were harvested by trypsinization and were treated with KCl (75mM) for 30min, followed by 10min treatment in fixative (1 part acetic acid and 3 part methanol). Cells were then spread on cold glass slides by dropping method followed by continuous flush with 1ml fixative for two times. Slides were air dried and then mounted with mounting medium containing DAPI (Vectashield). The mounted slides were kept in dark for 15 minutes. Imaging was performed on a fluorescence microscope (Axio observer.Z1, Carl Zeiss Micro-Imaging, Germany) attached with Axiocam HRM CCD camera and Apotome.2. Axiovision software (Zenpro2012).

## Author’s contributions

JC and BSK performed the experiments. KM and SG studied the FA mutations. RBM and SKM helped the iron uptake experiment. SSM planed the project and made the manuscript.

## Acknowledgments

We thank Dr. Agata Smogorzewska for providing the FA cell lines. Thanks to Jeffrey M. Rosen, Giovanni Pagano, K. Aikat, A. Bhattacharya for critical reading of the manuscript. This work was supported by Department of Biotechnology, Govt. of India. NIT Durgapur supported the fellowship to JC Bose K and BSK. Authors are thankful to DST-FIST for instrument grant to the Department of Biotechnology, NIT Durgapur. This work does not have any financial conflict of interest.

## Supplementary Materials

**Fig. S1.**
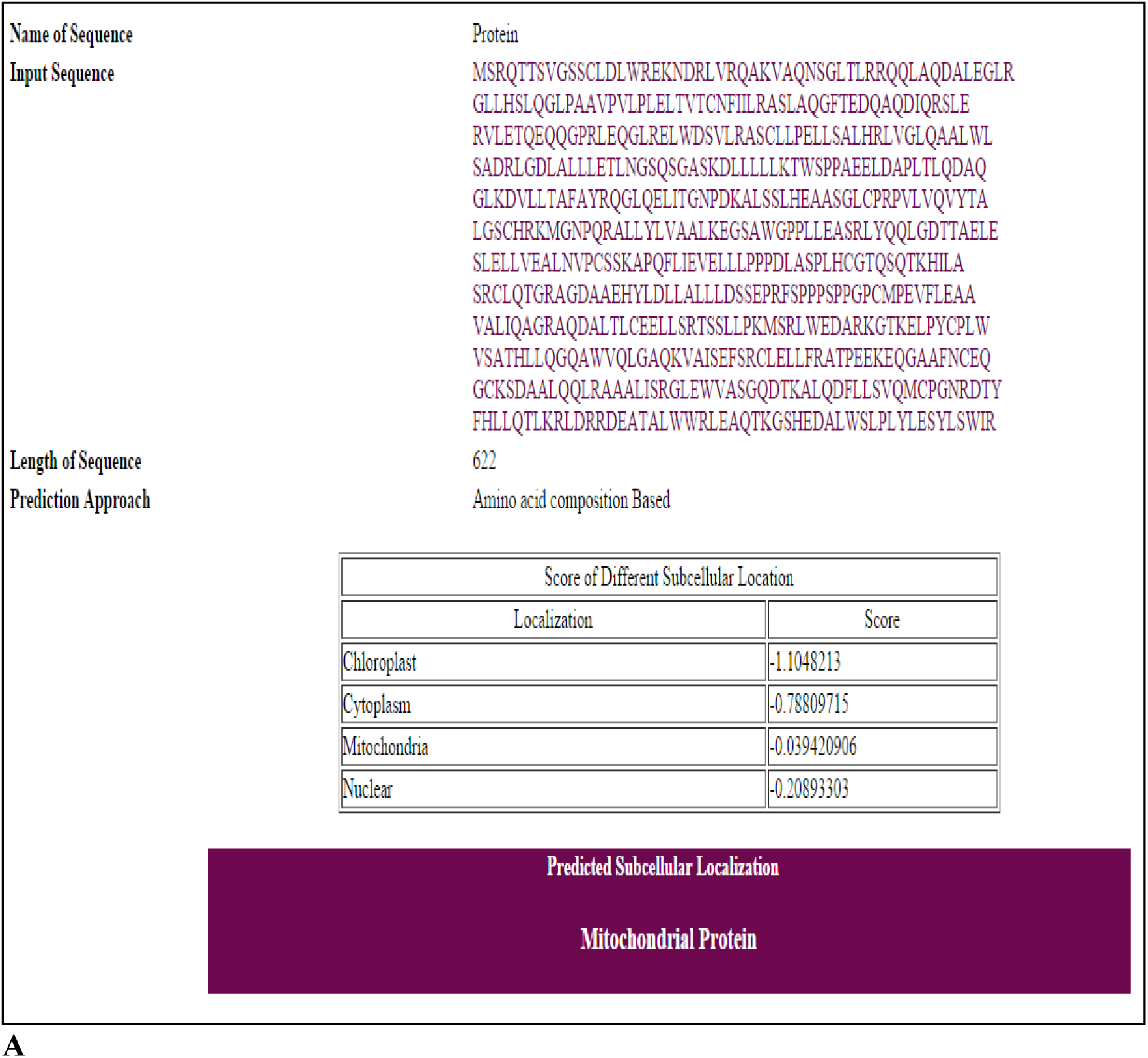
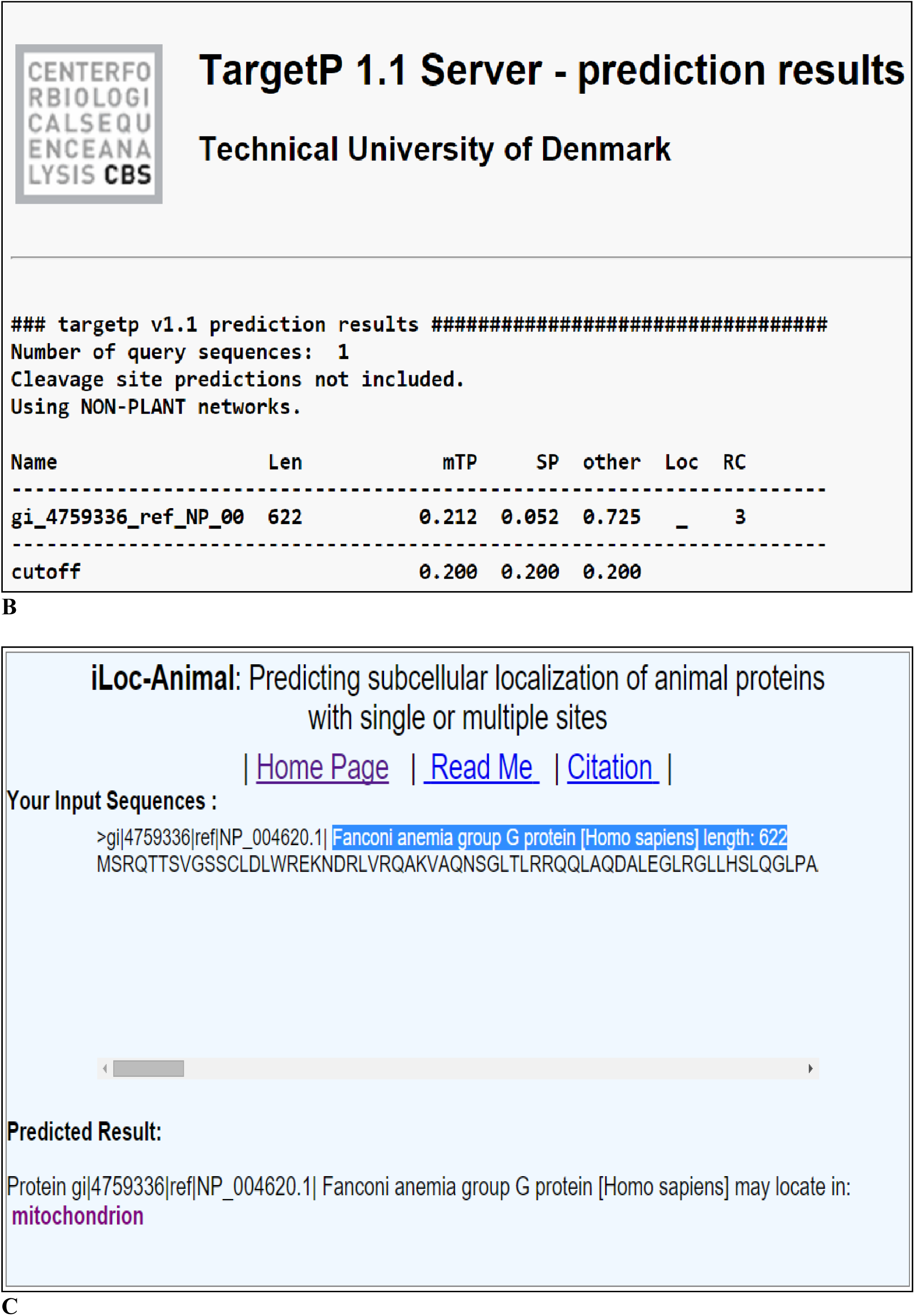
Insilico analysis of human FANCG for mitochondrial localization. **(A)**RSLpred score (cut off>-0.394209), **(B)** TargetP1.1 server score (0.212< cut off) and (**C)** iLOC-Animal results suggest mitochondrial localization.

**Fig. S2.**
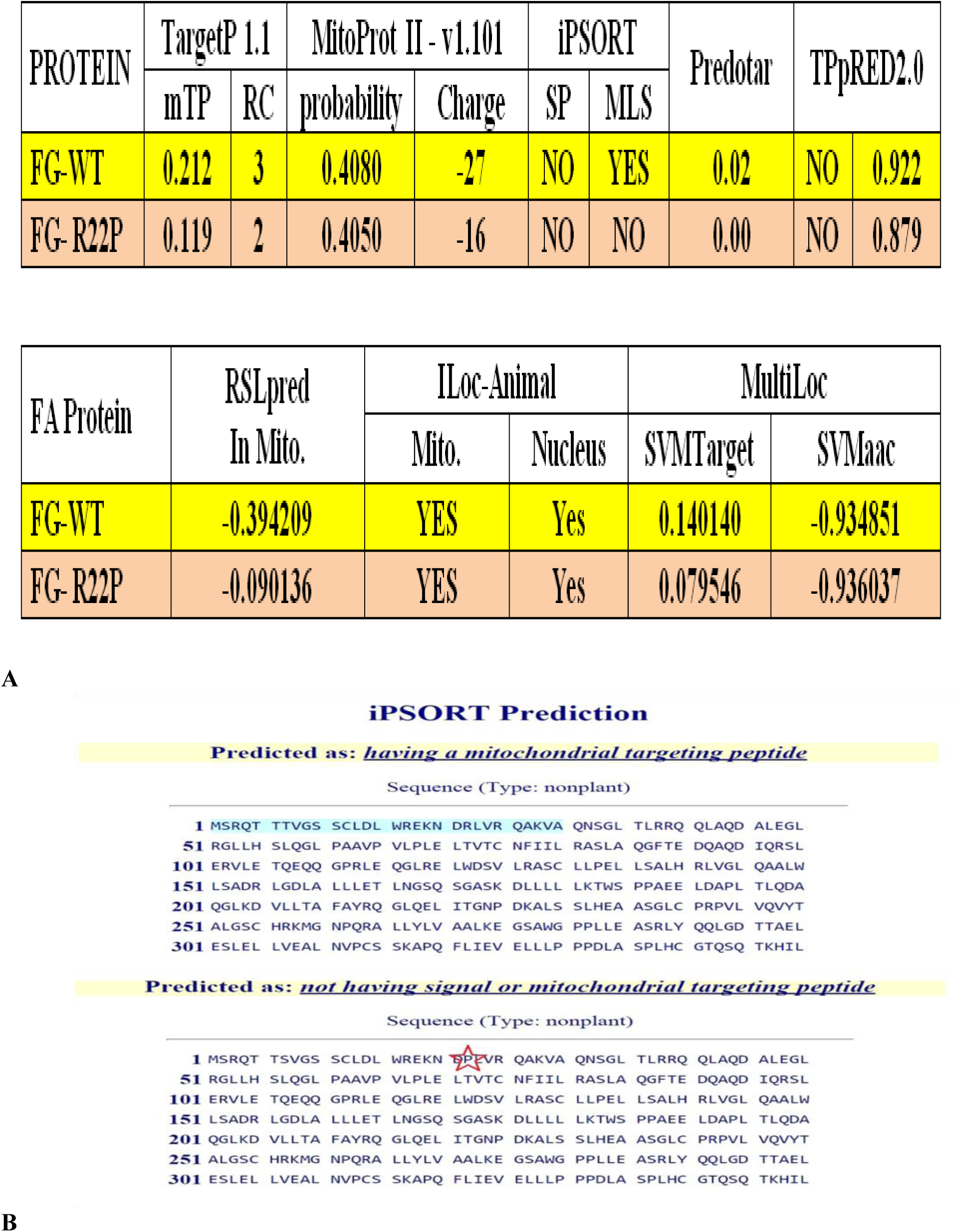
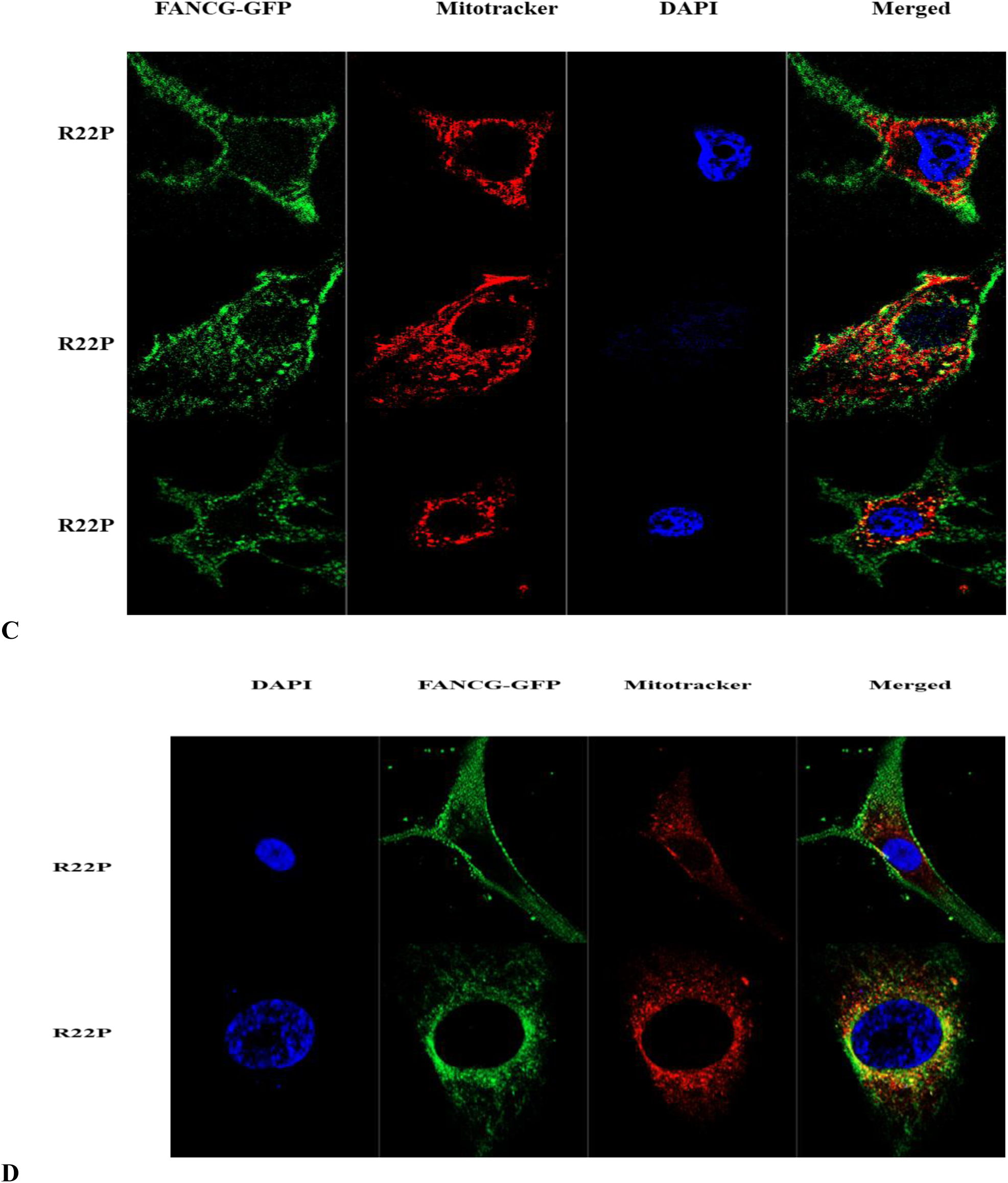
Mitochondrial localization of human FANCG-R22P mutant (FG-R22P) protein. The insilico tools **(A)** TargetP1.1, MitoProtII-v1.101, iPSORT, Predotar, TPpRED2.0, RSLpred, iLOC-Animal and MultiLoc have been used. mTP: mitochondrial targeting peptide; RC: Reliability class, SP: Signal peptide; MLS: Mitochondrial localization signal. **(B)** iPSORT analysis of FANCG (FG-WT) and FANCGR22P (FG-R22P) mutant protein for mitochondrial localization. **(C)** FG-R22P construct fused with GFP and Mitotracker have been transiently co-transfected into HeLa cells of different passages. and **(D)** FANCG parental cells. Co-localization of both GFP and Mitotracker has been observed (merged). Nucleus stained with DAPI.

**Fig. S3.**
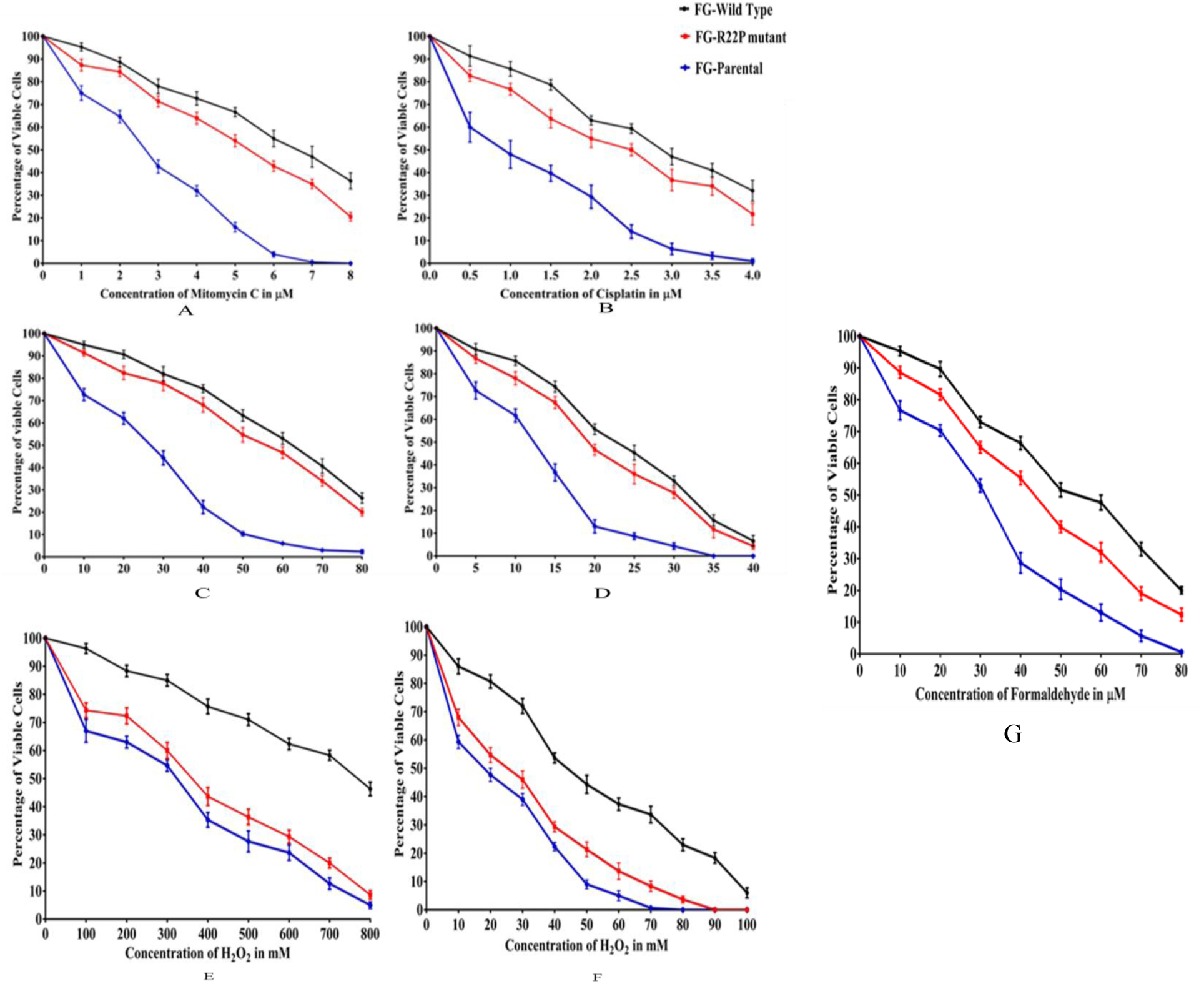
Drug sensitivity studies of FANCG corrected (black), FANCR22P (red) and FANCG parental cells (blue) Cells were treated with increasing concentration of drug (MMC and cisplatin) **(A & B)** for two days and **(C & D)** Five days, hydrogen peroxide (H2O2) for **(E)** two hrs and **(F)** twenty four hrs. **(G)**Formaldehyde for two hrs. Cell survivals were determined by Trypan blue assay. Each value is the mean of repeated (three times) experiments.

**Fig. S4.**
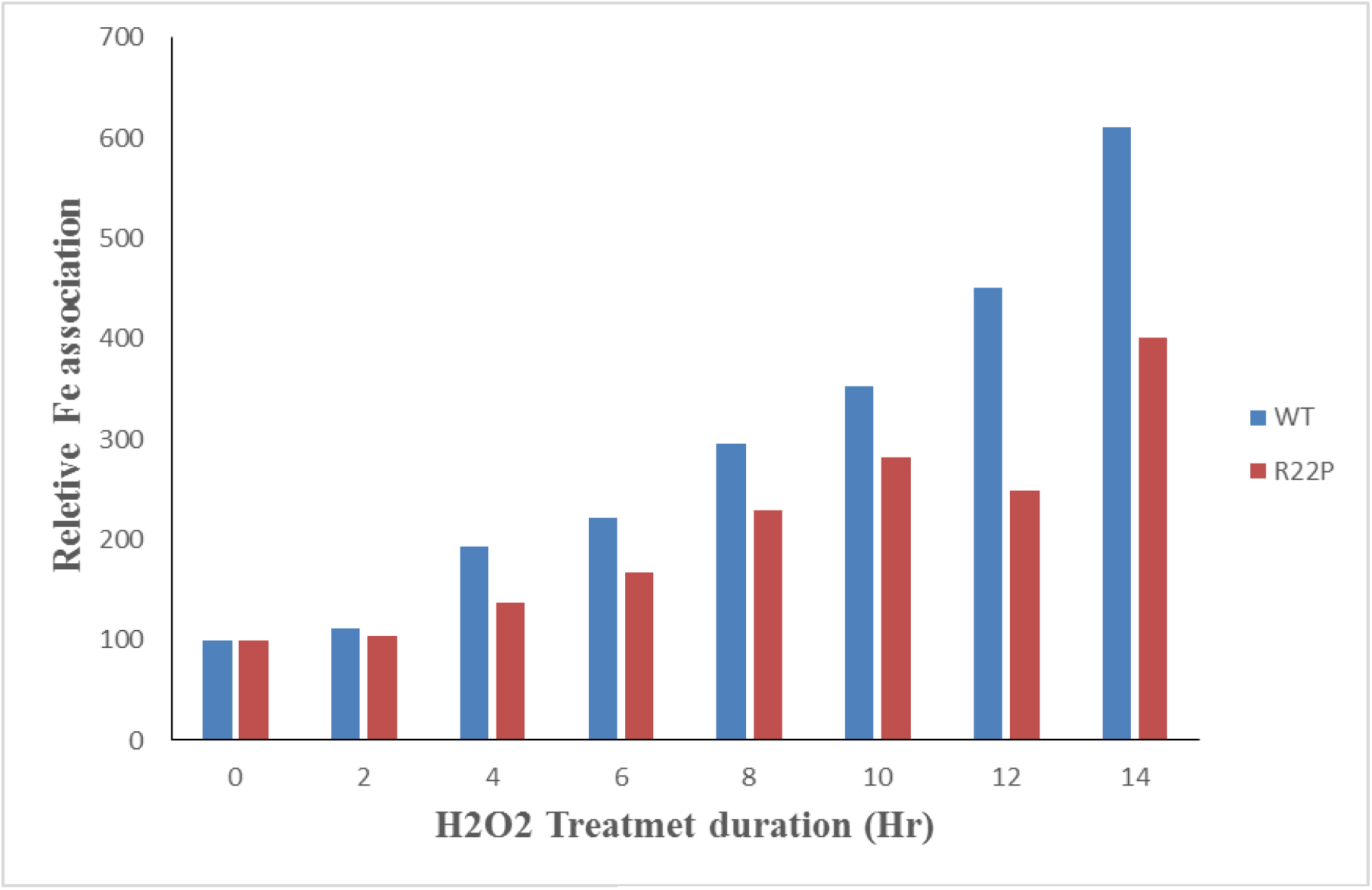
Fe association with Ferritin. Cells were incubated in IMDM media, containing ^55^Fe and treated with H_2_O_2_ (10µM) for fourteen hrs followed by MMC (100nM) treatment for thirty minutes at two hr intervals. Ferritin was immuno-precipitated (IP) using ferritin antibody (Sigma) and the concentration of ^55^Fe was determined. Concentration of ^55^Fe at 0 hr was considered as hundred. Blue bar reperesents FancG wild type fibroblast cells, Red bar represents FancG mutant (R22P) fibroblast cells.

**Fig. S5.**
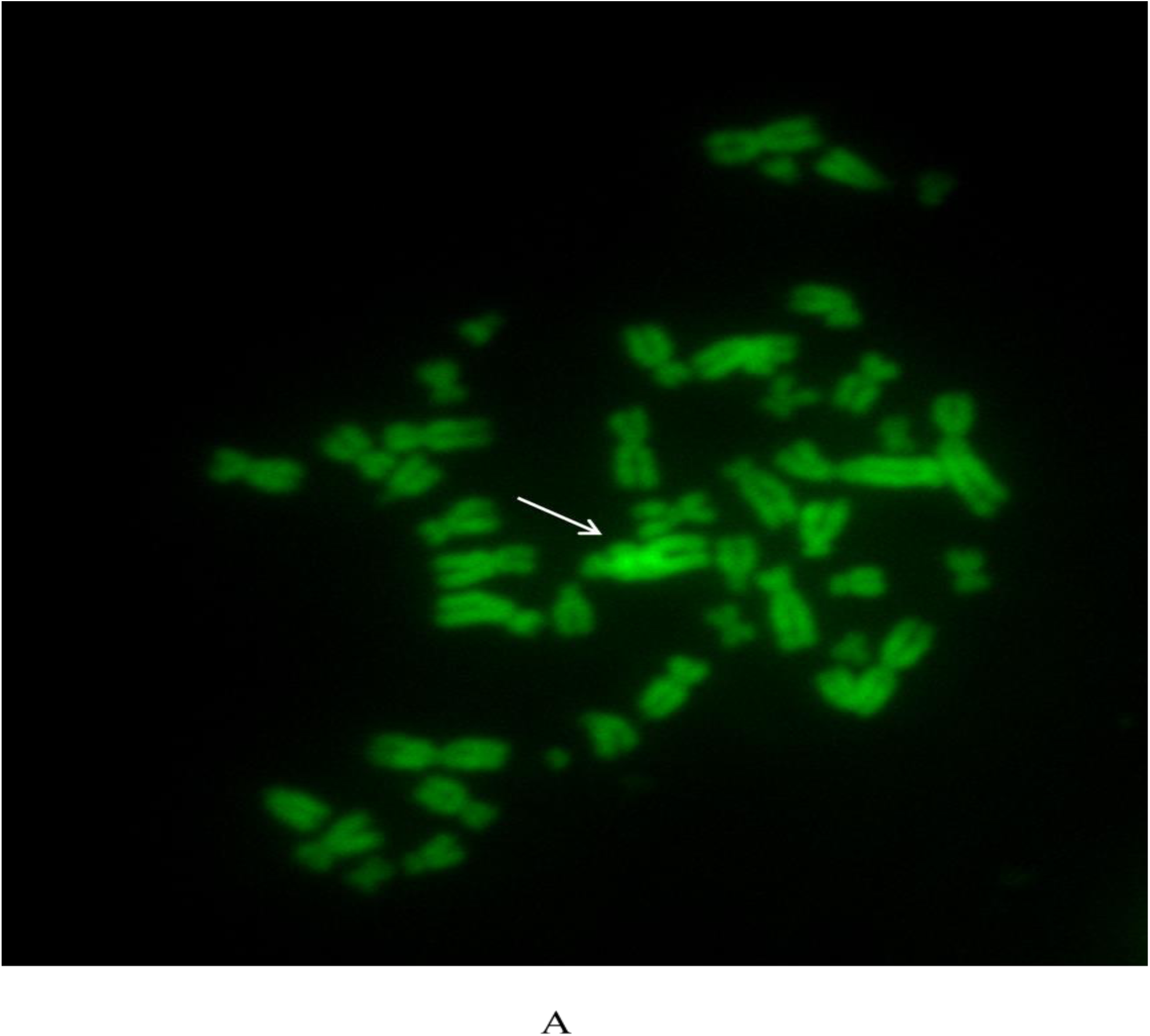
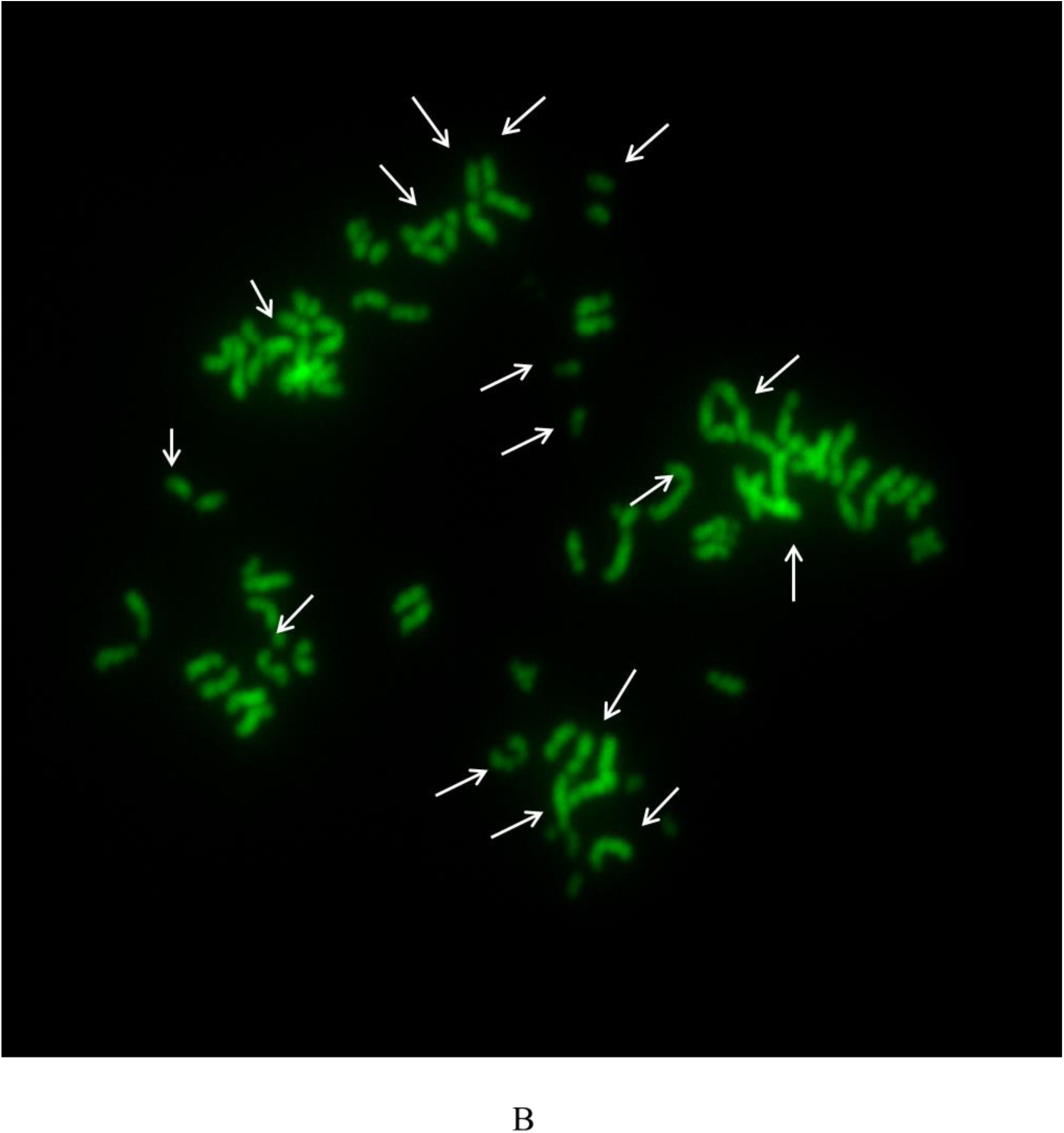
Chromosome preparation of R22P cells treated (A) with MMC and (B) with both MMC and H2O2. The arrows represent the deformed structure of the chromosome.

**Table.S1A.**

| PROTEIN | TargetP 1.1 |  | MitoProt II - v1.101 |  | iPSORT |  | Predotar | TPpRED2.0 |  |
| --- | --- | --- | --- | --- | --- | --- | --- | --- | --- |
|  | mTP | RC | probability | Charge | SP | MLS |  |  |  |
| FANCA | 0.223 | 3 | 0.0897 | -15 | NO | NO | 0.00 | NO | 0.985 |
| FANCB | 0.061 | 1 | 0.0732 | -05 | NO | NO | 0.00 | NO | 0.998 |
| FANCC | 0.059 | 2 | 0.0590 | -15 | NO | NO | 0.00 | NO | 0.995 |
| FANCD1 | 0.115 | 2 | 0.0169 | -30 | NO | NO | 0.01 | NO | 0.994 |
| FANCD2 | 0.182 | 2 | 0.2296 | -42 | NO | YES | 0.03 | NO | 0.999 |
| FANCE | 0.090 | 1 | 0.0070 | -20 | NO | NO | 0.00 | NO | 0.995 |
| FANCF | 0.056 | 2 | 0.1802 | +06 | NO | NO | 0.00 | NO | 0.725 |
| FANCG | 0.212 | 3 | 0.4097 | -13 | NO | YES | 0.02 | NO | 0.922 |
| FANCI | 0.054 | 1 | 0.1082 | -03 | NO | NO | 0.00 | NO | 0.993 |
| FANCJ | 0.093 | 2 | 0.0758 | -11 | NO | NO | 0.00 | NO | 0.987 |
| FANCL | 0.321 | 4 | 0.1797 | -05 | NO | NO | 0.02 | NO | 0.857 |
| FANCM | 0.875 | 2 | 0.5774 | -41 | NO | NO | 0.29 | YES | 0.977 |
| FANCN | 0.088 | 1 | 0.0090 | -01 | NO | NO | 0.00 | NO | 1.000 |
| FANCO | 0.577 | 4 | 0.3127 | -02 | NO | NO | 0.20 | NO | 0.898 |
| FANCP | 0.061 | 2 | 0.0081 | -27 | NO | NO | 0.00 | NO | 0.930 |
| FANCQ | 0.050 | 3 | 0.0699 | -07 | NO | NO | 0.00 | NO | 0.999 |
| FANCR | 0.059 | 1 | 0.0159 | -13 | NO | NO | 0.00 | NO | 0.997 |
| FANCS | 0.087 | 2 | 0.0248 | -86 | NO | NO | 0.00 | NO | 0.986 |
| FANCT | 0.346 | 4 | 0.2180 | 5 | NO | NO | 0.04 | NO | 0.984 |

**Table.S1.**
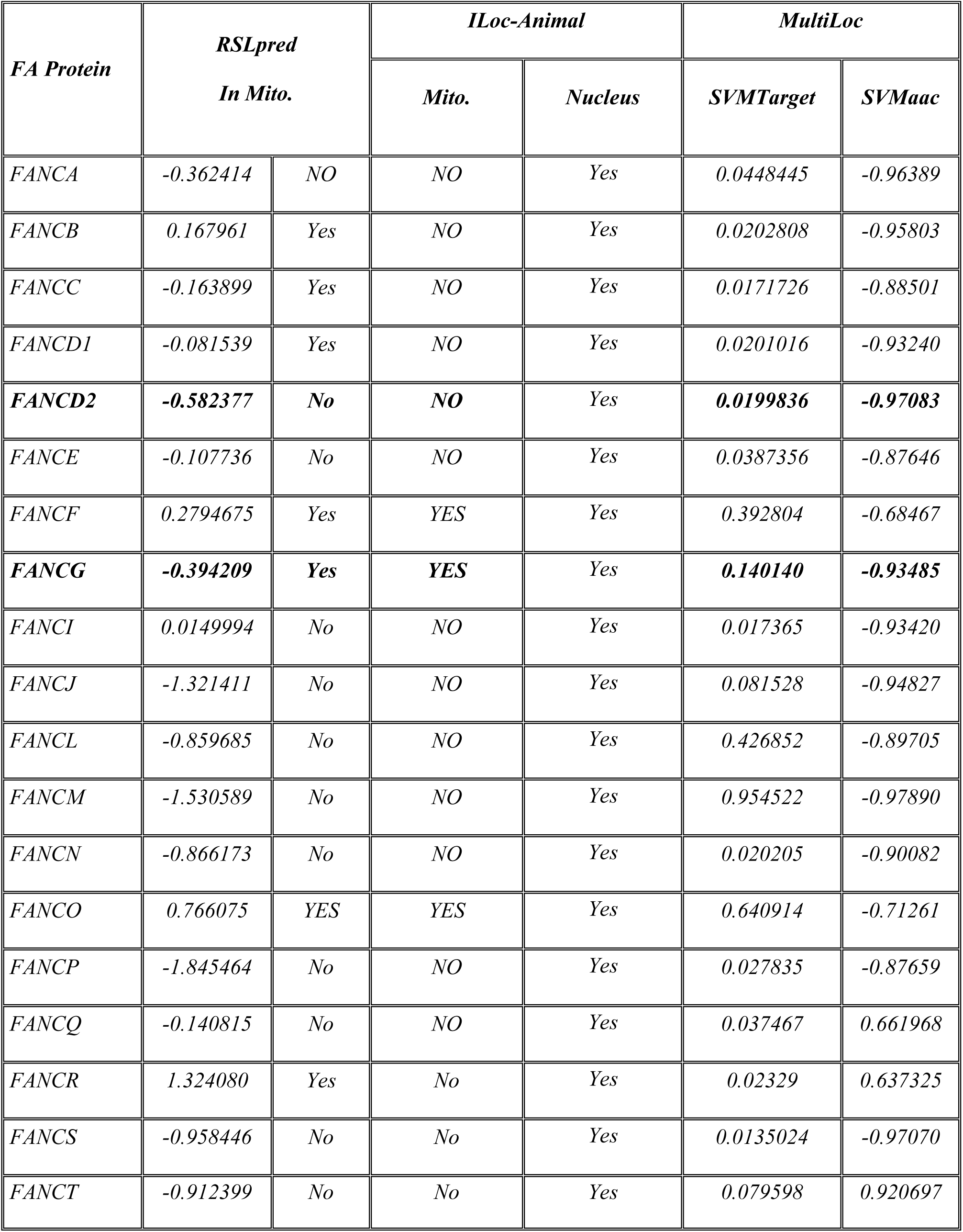
In silico analyses of Fanconi proteins for their cellular localization. The tools **(A)** TargetP1.1, MitoProt II –v1.101, iPSORT, Predotar, TPrRED2.0 **(B)** RSLpred,ILoc-Animal and MultiLoc have been used. Maximum probability of mitochondrial localization of FA proteins is shown in bold. mTP: mitochondrial targeting peptide; RC: Reliability class, SP: Signal peptide; MLS: Mitochondrial localization signal.

